# Heparin protects human neural progenitor cells from Zika Virus-induced cell death and preserves their differentiation into mature neural-glia cells

**DOI:** 10.1101/2021.05.05.442746

**Authors:** Isabel Pagani, Linda Ottoboni, Paola Podini, Silvia Ghezzi, Elena Brambilla, Svetlana Bezukladova, Davide Corti, Marco Emilio Bianchi, Maria Rosaria Capobianchi, Edwin A Yates, Gianvito Martino, Elisa Vicenzi

## Abstract

Zika virus (ZIKV) is an arbovirus member of the Flaviviridae family that causes severe congenital brain anomalies in infected fetuses. Human neural progenitor cells (hNPCs) are highly permissive to ZIKV infection, causing inhibition of cell proliferation concomitant with an induction of cell death. We previously demonstrated that pharmaceutical-grade heparin inhibited virus-induced cell death with minor effects on *in vitro* virus replication in ZIKV-infected hNPCs. Here we show that heparin prevented ZIKV-induced intracellular vacuoles, a signature characteristic of paraptosis, but also inhibited necrosis and apoptosis of hNPCs when grown as neurospheres (NS). Furthermore, heparin preserved the differentiation of both ZIKV-infected human-induced pluripotent stem cells (hiPSC) derived-NPCs and fetus-derived NPCs into neural-glial cells. Collectively, these results highlight the potential neuroprotective effect of heparin that could be re-purposed and exploited to drive the development of novel agents for preventing ZIKV damage.

## INTRODUCTION

Zika virus (ZIKV) is a member of the *Flaviviridae* family that is mainly transmitted to humans through the mosquito bites of the *Aedes* species ^1^, but it can also spread through sexual transmission ^2, 3^, maternal-fetal pathway, and blood transfusion ^4, 5^. After its discovery in 1947 ^6^, ZIKV human infections were occasionally reported in Africa and Asia, typically with mild clinical presentation ^7^. After 70 years of limited ZIKV spread with a few outbreaks such as those in the Pacific Islands in 2007 ^8^ and French Polynesia in 2013 ^9^, a significant challenge emerged to global health authorities in February 2016 following the unexpected increase of neonates affected by abnormally small brain (microcephaly) in South America ^10, 11^. The widespread ZIKV infection of pregnant women had caused serious birth defects, including neurological diseases ^12, 13, 14^. *In utero* ZIKV-associated pathological conditions of the fetus were supported by the presence of ZIKV particles visible in brain sections by transmission electron microscopy and recovery of the full ZIKV genome from the fetal brain ^15^. By immunolabelling of brain from fetus with ZIKV intrauterine infection, ZIKV was detected in neural and non-neural cells, although the highest rate of infection was in intermediate progenitor cells and immature neurons ^16^, suggesting a strong tropism for human neural progenitor cells (hNPCs). Numerous reports demonstrated that human pluripotent stem cell (hPSC)-induced NPCs are highly permissive to ZIKV *in vitro* infection and that this causes inhibition of their proliferation with concomitant cell death ^17,18, 19, 20, 21, 22, 23, 24^. These findings were also reproduced in mouse animal models which further demonstrated the development of microcephaly in infected fetal mice ^25, 26, 27^.

Several studies have focused on the modalities of ZIKV-induced cell death and both apoptotic and non-apoptotic modalities have been described as mechanisms that lead hNPCs to die during viral replication ^28, 29^. ZIKV, like other flaviviruses, exploits the endoplasmic reticulum (ER) to assemble the replication complex with an expansion of cellular membranes that contain newly formed immature virions ^30^. This overwhelming ER overload leads to ER stress with consequent stimulation of the unfolded protein response ^31^ and, in particular, expression of the C/EBP homologous protein (CHOP) that initiates apoptosis in infected cells ^32^. Prolonged ER stress upon ZIKV infection, however, triggers a particular type of non-apoptotic cell death characterized by the cytoplasmic formation of ER-derived vacuoles which are formed in so called paraptosis-like cell death ^33^, first defined with morphological criteria and independently of the activation of cleaved-caspase 3 (cl-CASP3) ^34^. A hallmark of non-apoptotic cell death is the expression of high mobility group 1 (HMGB1) protein ^35, 36^. HMGB1 was first described as a DNA binding protein ^37^, however, while in the nucleus it is involved in multiple functions, such as transcription, replication, recombination, DNA repair and genomic stability ^38^, upon cell death, HMGB1 can be passively released into the extracellular environment where it acts as a damage-associated molecular patterns (DAMP) to activate host defenses ^39^.

Following the strategy of repurposing drugs already approved for another medical need, we described the ability of heparin to uncouple viral replication from virus-induced cell death by inhibiting ZIKV cytopathic effect without affecting viral replication ^40^. In the present study, we have tested whether heparin can prevent ZIKV-induced apoptotic and non-apoptotic/paraptosis cell death in the infection of stem-cell-derived neural progenitors. As hiPSC-derived NPCs do not consistently recapitulate *in vivo* embryonic human brain gene expression ^41^, we employed human fetus(hf)-derived NPCs obtained directly from the tissue that is the main target of ZIKV infection ^16, 23^. In addition, both hiPSC-NPCs and hf-NPCs aggregate to form 3D neurospheres (NS) when grown in non-adherent conditions and have been used as a model of *in vivo* neurogenesis phenotype for ZIKV infection ^18, 22^. We produced NS from both hiPSC-NPCs and hf-NPCs to determine the effect of heparin protection on ZIKV infection.

As heparin prevented virus-induced cell death with partial reduction of viral replication, we tested whether in ZIKV-infected hNPCs, heparin preserved the ability of hNPCs to differentiate into mature neural-glial cells. To this end, both hiPSC-NPCs and hf-NPCs were infected with ZIKV and then differentiated into neuro-glial cells. Heparin was evaluated prior to, or after, infection and was repeatedly added during the differentiation experiment.

Our results highlight the potential to delay ZIKV infection and replication and indicate that heparin is endowed with neuro-protective activity against the cytopathic effect of ZIKV in hNPCs.

## RESULTS

### Heparin inhibits ZIKV-induced non-apoptotic cell death

We have previously demonstrated that heparin prevents ZIKV-induced cells death in hNPCs grown as monolayers ^40^. Since a recent report highlighted a mechanism of massive cytoplasmic vacuolization that leads to paraptosis-like death in ZIKV-infected HeLa cells, human foreskin fibroblasts and astrocytes ^33^, we tested whether vacuoles were present in infected hNPCs derived from reprogramming human induced-pluripotent stem cells (hiPSC-NPCs) and whether heparin affected the vacuole formation. Firstly, cells infected with the PRVABC59 isolate were examined at different time points post-infection by transmission electron microscopy (**Fig. 1**). Single virions were visible in the cytoplasm and in the lysosomes shortly after infection (4 h) and tended to aggregate and then localized around membrane-delimited vacuoles 9 and 16 h post-infection. Occasionally, virions were contained inside the vacuoles, but the majority was located outside. Heparin treatment 1 h prior infection (100 µg/ml) did not alter the virion density or location compared to untreated cultures until 16 h post-infection (**Fig. 1A**). The number of vacuoles increased 72 h post-infection in the infected untreated cultures, whereas heparin reduced virion-associated vacuoles and cells exhibited healthy and normal-appearing mitochondria (**Fig. 1B**). In order to quantify the effect of heparin on ZIKV-induced vacuoles, virion positive cells were examined for the presence of vacuoles. A total of 217 out of 247 ZIKV infected cells were vacuole positive (88%, **Fig. 1C**). In contrast, heparin significantly reduced the number of vacuole-positive cells to 52 out of 229 (23%). As membrane damage causes the passive release of HMGB1 protein, a marker of non-apoptotic cell death, HMGB1 release was measured in the culture supernatant of ZIKV infected hiPSC-NPCs in the presence or absence of heparin. The kinetics of HMGB1 release showed that HMGB1 was present at low levels in the cell culture supernatant of uninfected hiPSC-NPCs(**Fig. 1D**); however, ZIKV infection significantly increased the levels of HMGB1 after 6 days of infection while heparin significantly lowered HMGB1 levels to those of uninfected controls (**Fig. 1D**).

**Fig. 1.**
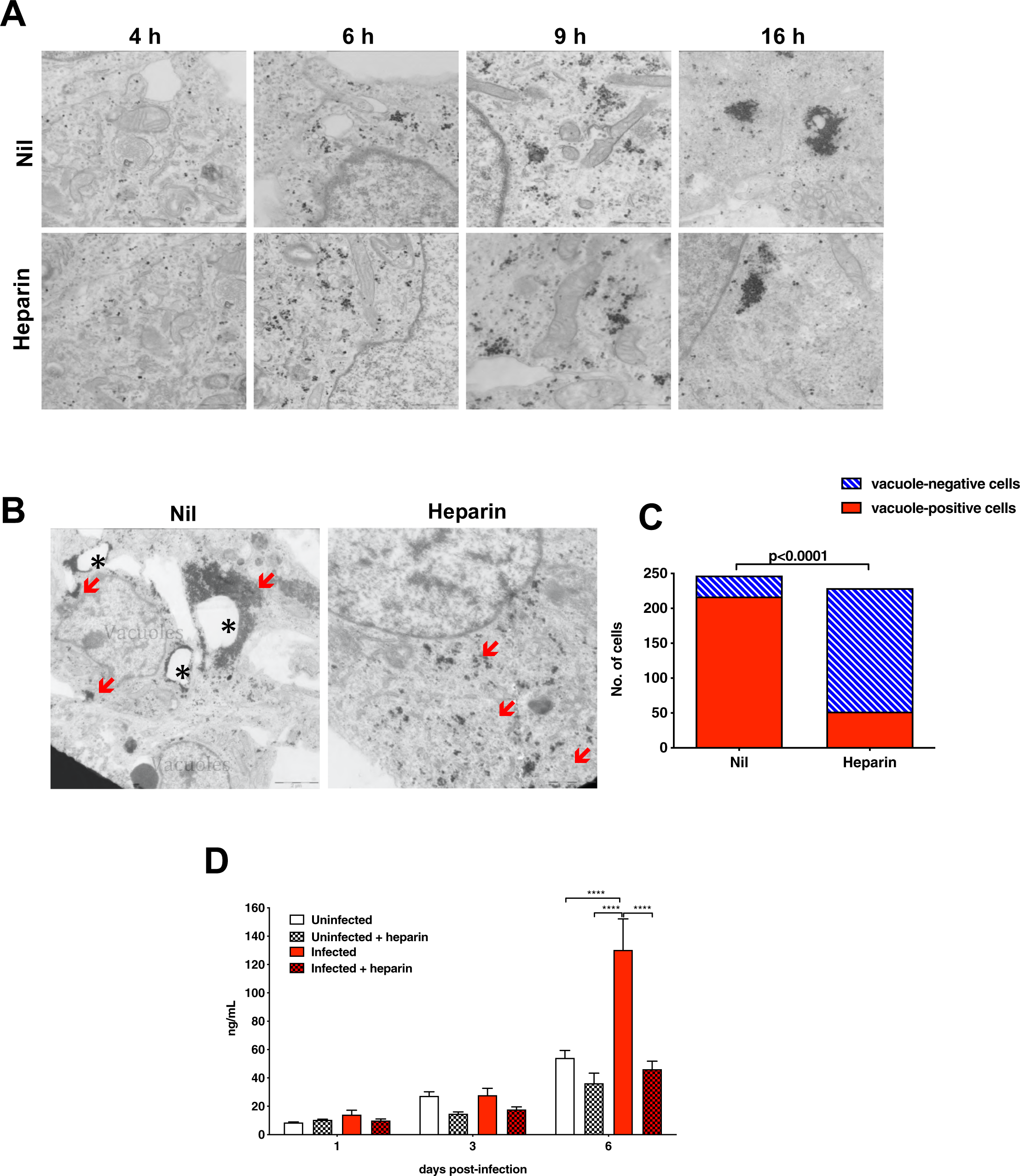
Heparin prevents the formation of cytoplasmic vacuoles in ZIKV-infected hiPSC-derived NPCs. **A.** Electron microscopy specimens of hiPSC-NPCs treated, or not, with heparin (100 µg/ml) 1 h prior to infection with the PRVABC59 isolate (MOI of 1). Infected hiPSC-NPCs were fixed at 4, 6, 9, and 16 h post-infection. Increased accumulation of clusters of ZIKV particles were visible in untreated and treated infected hiPSC-NPCs; bar = 2 µm. **B.** Electron microscopy specimens of hiPSC-NPCs treated, or not, with heparin (100 µg/ml) 3 days post-infection. Asterisks (*) indicate vacuoles and clusters of ZIKV particles are indicated with red arrows; bar = 2 µm. **C.** Quantification of vacuoles in hiPSC-NPCs treated, or not, with heparin prior to infection with infected with the PRVABC59 isolate. Bars represent vacuoles-positive (red) and vacuoles-negative (blue) cells counted in more than 200 images taken from 3 independent experiments. p<0.0001 Fisher’s extract test. **D.** Levels of HMGB1 released in the culture supernatants after 1, 3 and 6 days post-infection. Bars represent the mean ± SD of 3 independent experiments. Two-way Anova with the Bonferroni correction was used. * Represents statistical comparison among groups (****, p<0.0001).

Taken together these results suggest that the ZIKV-induced cytopathic effect from non-apoptotic, paraptosis-like damage was reverted by heparin treatment.

### Heparin inhibits necrosis of ZIKV-infected neurospheres (NS)

In order to validate heparin protection of ZIKV-induced cell damage observed in hiPSC-NPCs grown as a monolayer, a more complex 3D system of NS formed by NPC aggregates ^42^ was infected with ZIKV in the presence or absence of exogenous heparin. In addition to hiPSC-NS, NS were obtained from hNPCs isolated from a human fetal brain (hf-NS). First, to determine the efficiency of ZIKV infection in hf-NPCs as compared to hiPSC-NPCs, cells in monolayer were infected with three ZIKV isolates including the original African MR766 and two recent ones, i.e. the Puerto Rico PRVABC59-2015 and the Brazilian 2016-INMI-1. As shown in **Fig. 1S** **A**, the peak of viral replication was reached at day 3 post-infection in both cell systems. In hiPSC-NPCs, a decrease of infectious virus released in the supernatant was detected at day 10 post-infection being more prominent with the infection of MR766 compared to the two more recent isolates (**Fig. 1S** **A**). In hf-NPCs, infectious virus persisted up to 10 days post-infection (**Fig. 1S B**) consistently with lower levels of cytopathicity as measured by the adenylate kinase (AK) activity released in the supernatant of hf-NPCs as compared with hiPSC-NPCs (**Fig. 1S D** and **C**, respectively). The contemporary Brazilian 2016-INMI-1 isolate was selected for the next NS infection experiments.

NS, spontaneously formed from hNPCs after 3 days of culture in rotation without coating of extracellular matrix, were treated with heparin (100 µg/ml) 1 h prior to infection with the Brazilian 2016-INMI-1 isolate. The number and diameter of hiPSC-NS was evaluated 3 days post-infection. ZIKV infection caused a marked reduction of hiPSC-NS number and size, whereas heparin reverted this effect almost to the levels of uninfected NS (**Fig. 2A**). Indeed, the diameter of uninfected hiPSC-NS had an average size of 513±188 μm that was not statistically different from that of heparin-treated uninfected NS, whereas ZIKV-infected NS had a significantly smaller average size (241±80 µm). Interestingly, heparin preserved the diameter of the infected NS to the levels of uninfected ones (449±130 µm) (**Fig. 2B**). In order to determine whether the size reduction was dependent on virus-induced cell death, the extent of ZIKV-induced necrosis was determined. The reduction of number and size of infected NS was consistent with the levels of necrosis measured by the AK activity in the culture supernatant. As shown in **Fig. 2C**, the amount of AK activity released in the culture supernatant significantly increased in infected cultures, whereas heparin decreased the AK activity levels to those of control cultures.

**Fig. 2.**
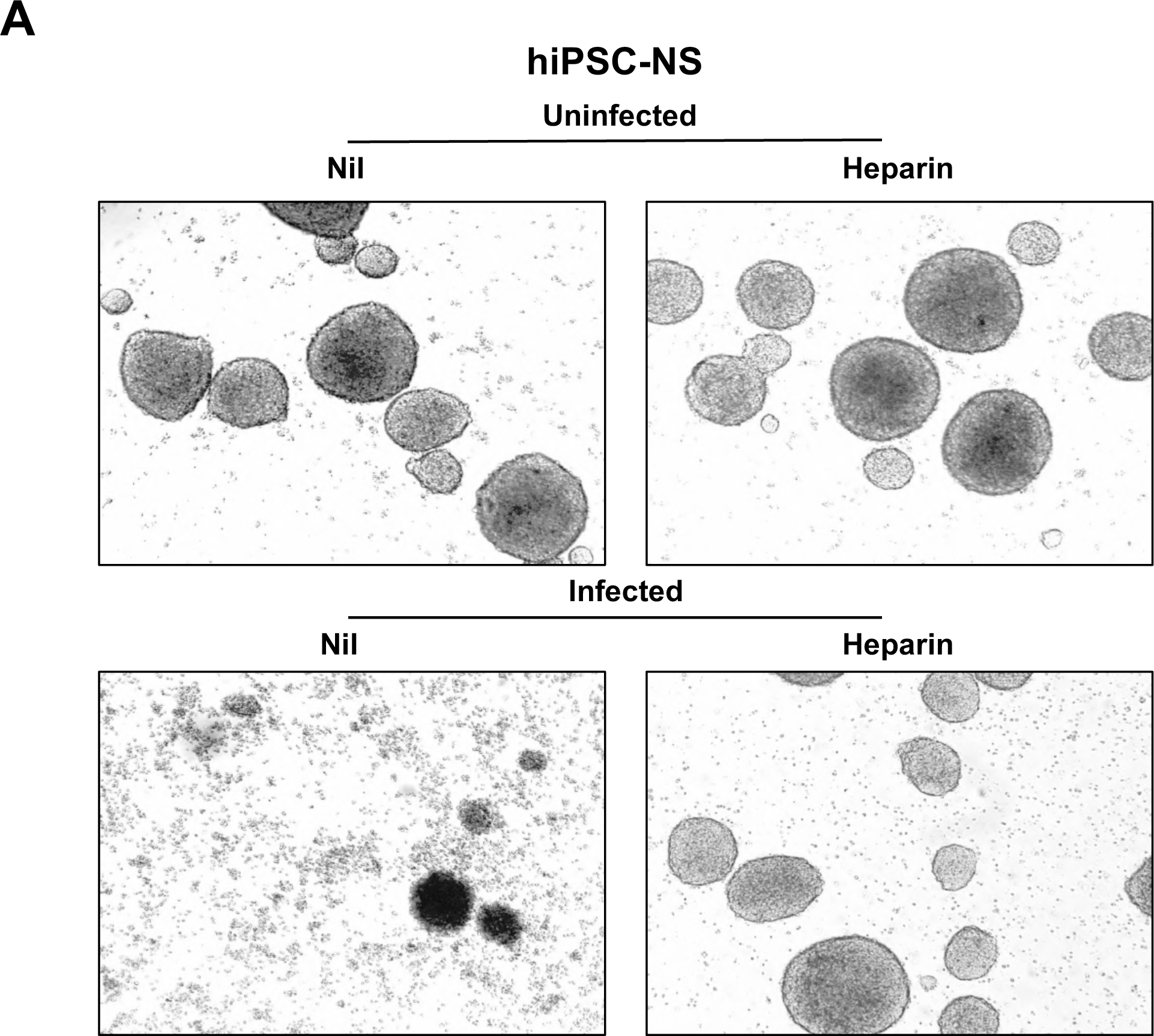
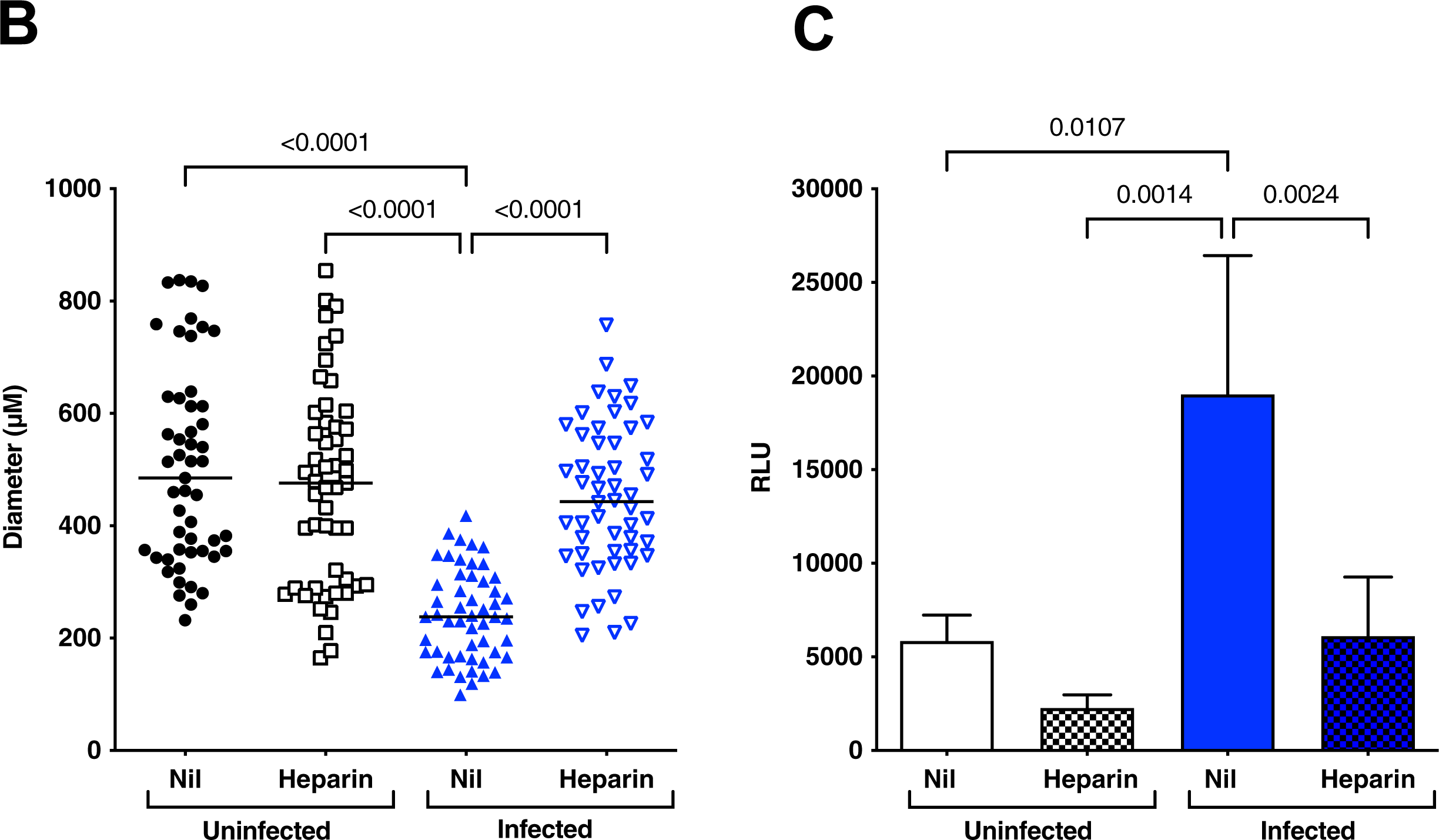
Heparin inhibits disruption of ZIKV-infected hiPSC-derived NS. **A.** hiPSC-NS were treated with heparin (100 μg/ml) 1 h prior to infection with Brazilian 2016-INMI-1 isolate (MOI of 1). After 4 h of ZIKV adsorption, virus inoculum was removed and the medium was replenished. Bright field photomicrographs of mock-treated *vs.* heparin in both uninfected and infected conditions were taken 6 days post-infection. Calibration bars: 150 μm. **B.** ZIKV caused a significant reduction in the NS number and size that was reverted by heparin to the levels of control cultures. Individual NS diameter values with the mean ± SD are shown. P values were calculated by one-way Anova with the Bonferroni correction. **C.** Supernatants were collected 6 days post-infection and tested for AK activity. The results are expressed as relative luminescent units (RLU). Bars represent the mean ± SD of 5 independent experiments. P values were calculated by one-way Anova with the Bonferroni correction.

Similarly, ZIKV infection induced a reduction of number and size in hf-NS (**Fig. 3A**). The diameter of uninfected cultures had an average size of 485±86 µm, whereas that of infected hf-NS was significantly reduced (185±56 µm). Heparin partially preserved the diameter of the hf-NS to that of the uninfected treated control (393±71 µm) (**Fig. 3B**). As shown in **Fig. 3C**, the amount of AK activity released in the culture supernatant significantly increased in infected cultures, whereas heparin decreased the AK activity to that of control cultures.

**Fig. 3.**
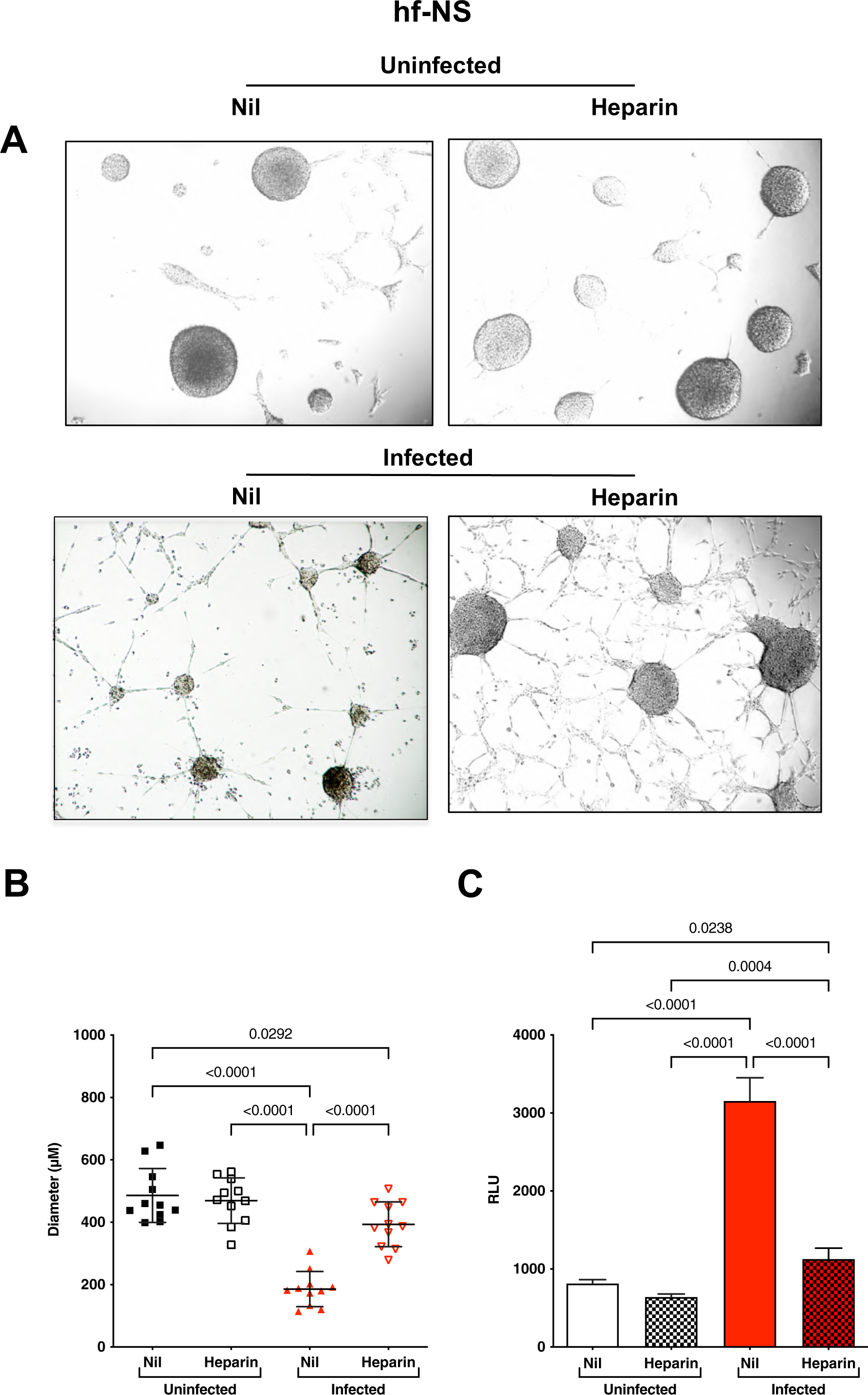
Heparin inhibits disruption of ZIKV-infected hf-NS. **A.** hf-NS were treated with heparin (100 μg/ml) 1 h prior to infection with the Brazilian 2016-INMI-1 isolate (MOI of 1). After 4 h of ZIKV adsorption, virus inoculum was removed and the medium was replenished. Bright field photomicrographs of mock-treated vs. heparin in both uninfected and infected conditions were taken 6 days post-infection. Calibration bars 150 μm. **B.** ZIKV caused a significant reduction in NS number and size that was reverted by heparin to the levels of control cultures. Individual NS diameter values with the mean ± SD are shown. P values were calculated by one-way Anova with the Bonferroni correction. **C.** Supernatant of infected hf-NS with the Brazilian 2016-INMI-1 isolate was collected 6 days post infection and tested for AK activity. The results are expressed as relative luminescent unit (RLU). Bars represent the mean ± SD of 3 independent experiments. One-way Anova was used with the Bonferroni correction.

### Heparin inhibits apoptosis of ZIKV-infected NS

Since heparin treatment protected NS from ZIKV-induced disruption, the architecture of hiPSC-NS was determined by immunofluorescence. As shown in **Fig. 4A**, vimentin staining, a type III intermediate filament of the cytoskeleton, formed a web inside uninfected NS. Six days after infection, however, infected hiPSC-NS lost their integrity and vimentin appeared like dots resembling agglomeration. Conversely, following heparin treatment the vimentin web was still maintained even in the presence of virions. As the maintenance of NS integrity could be due to a decrease of both overall NS damage and cell death induced by ZIKV infection, NS were stained with an anti-cl-CASP3 monoclonal antibody (mAb), to determine the level of apoptosis. As shown in **Fig. 4B**, the cl-CASP3 signal in the heparin-treated cells was lower than the untreated infected one. Notably, in uninfected NS, apoptosis was induced spontaneously in the core of NS as described previously ^43^.

**Fig. 4.**
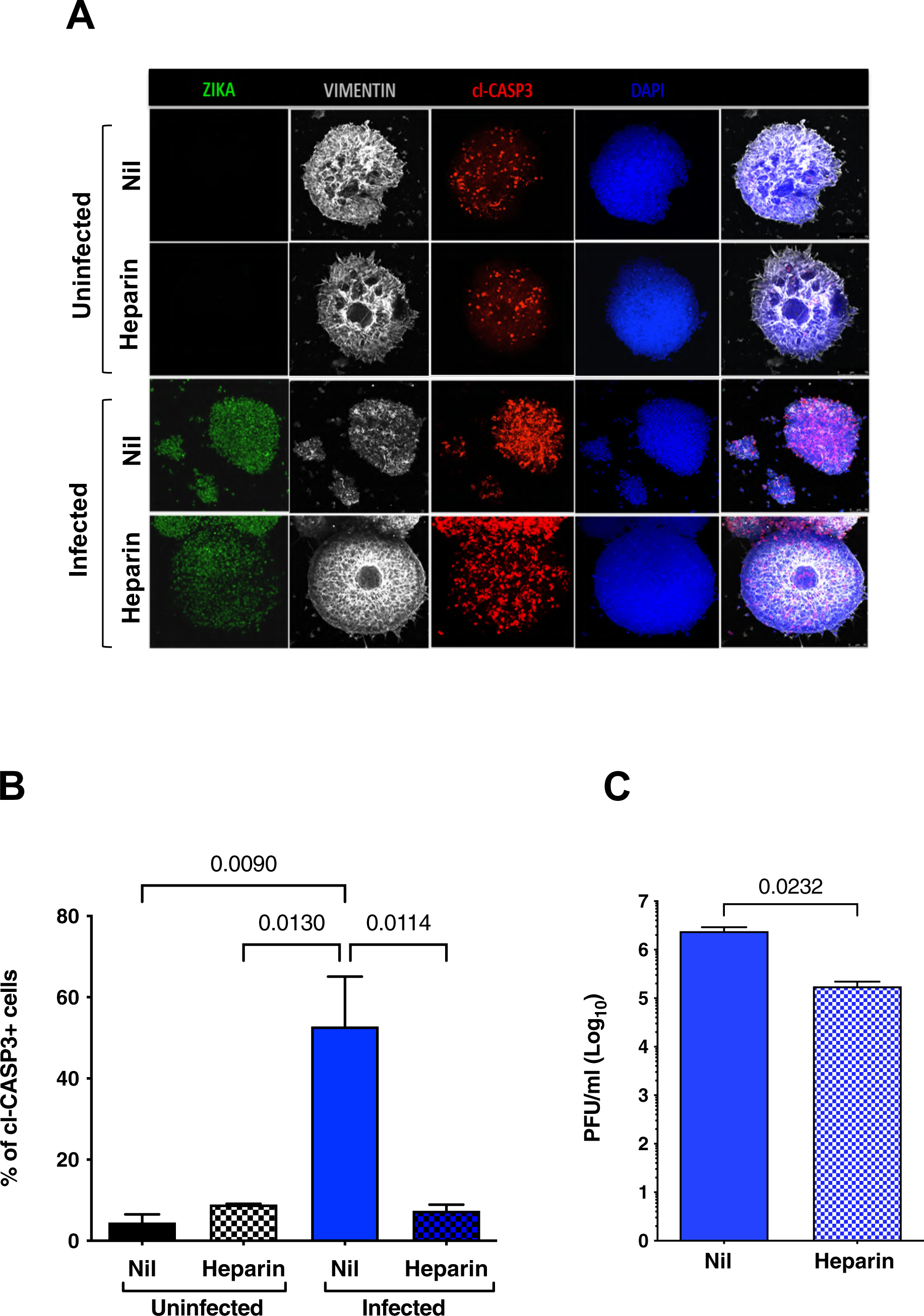
Heparin preserves hiPSC-NS structural integrity. **A.** hiPSC-NS were treated with heparin 1 h prior to infection with Brazilian 2016-INMI-1 isolate (MOI of 1). After 4 h of ZIKV adsorption, virus inoculum was removed and the medium was replenished. hiPSC-NS were fixed after 6 days post-infection. Immunostaining for dsRNA (green), vimentin (white) and cl-CASP3 (red) counterstained with DAPI (blue) in uninfected, ZIKV-infected and heparin-treated NS. Scale bar: 50 μm. Images were obtained using a SP8 confocal microscope. **B.** hiPSC-NS were fixed after 4 and 6 days post-infection and immunostained for cl-CASP3 antibody. Scale bar: 50 μm. Images were obtained using a SP8 confocal microscope. Z-stacks analyses were performed on 3 independent images for each condition and time points. **C.** Six days post-infection, infectious viral titers were determined in the supernatant by a plaque forming assay in Vero cells. Bar graphs indicate the plaque forming unit /ml (PFU/ml). Means ± SEM of 3 independent experiments are shown. P values were calculated by Student’s paired t-test.

As heparin prevented ZIKV-induced cytophatic effect, we tested whether the amount of infectious virus released into the culture supernatant was modified by heparin-treated NS. As shown in **Fig.4C**, the infectious titers were significantly lowered by approximately 10-fold in heparin-treated NS as compared to infected untreated NS, suggesting that heparin might also inhibit viral entry and replication, as previously reported for other viruses including SARS-CoV-2 ^44, 45^.

### Treatment of hiPSC-NPCs with heparin either before or after infection preserves differentiation into neural cells

As heparin inhibits ZIKV-infected hNPCs to undergo ZIKV-induced cytopathicity with virions inside the cells, our next goal was to test whether heparin preserved hiPSC-NPC differentiation into mature neural-glial cells ^46, 47, 48^. To this end, differentiation of hiPSC-NPCs was induced following a two-step approach modified from Muratore et al. ^49^ beginning after 4 h of incubation with ZIKV. After virus inoculum removal, hiPSC-NPCs were maintained in induction medium for 7 days, followed by a 14 day culture in differentiation medium ^49^. Cells were incubated with heparin (100 µg/ml) either 1 h prior to infection (Pre-treatment) or 1h after removal of the virus inoculum (Post-treatment). Indirect immunofluorescence was performed to detect infected cells by staining with a human mAb against ZIKV-E (envelope) protein in addition to an anti-Pax6 and anti β-III-tubulin (TUJ1) mAbs that stain neural progenitor cells and immature neurons, respectively. As shown in **Fig. 5A**, productive ZIKV infection was measured in untreated cultures with a peak of virus replication 8 days post-infection. Then, virus production started to decrease progressively due to a strong virus-induced cytopathic effect as shown by indirect immunofluorescence at days 12 and 14 post-infection (**Fig. 5B**). In contrast, when cells were incubated with heparin either pre- or post-infection, a 4-day delayed peak of virus replication was observed. Cultures incubated with heparin before infection released significantly lower levels of virions in the culture supernatant than controls as early as 1 day up to day 7 post-infection (**Fig. 5A**). As shown in **Fig. 5B**, the proportion of cells positive for viral antigens was lower in heparin-treated cells than in untreated cells, consistent with the inhibition of viral entry and replication. As expected, at the end of the differentiation period (day 14), staining of the cells with TUJI, a marker of neuron differentiation, showed that TUJI-positive cells were more visible in uninfected-untreated cultures compared to day 12 when Pax6 staining was still abundant (**Fig. 5B**). By preventing virus-induced cytopathic effect heparin partially preserved the differentiation of hiPSC-NPCs into neurons regardless of whether it was added before or after infection. This effect lasted at least up to day 12 post-infection and also resulted in delaying the virus-induced cytopathic effect (**Fig. 5B**).

**Fig. 5.**
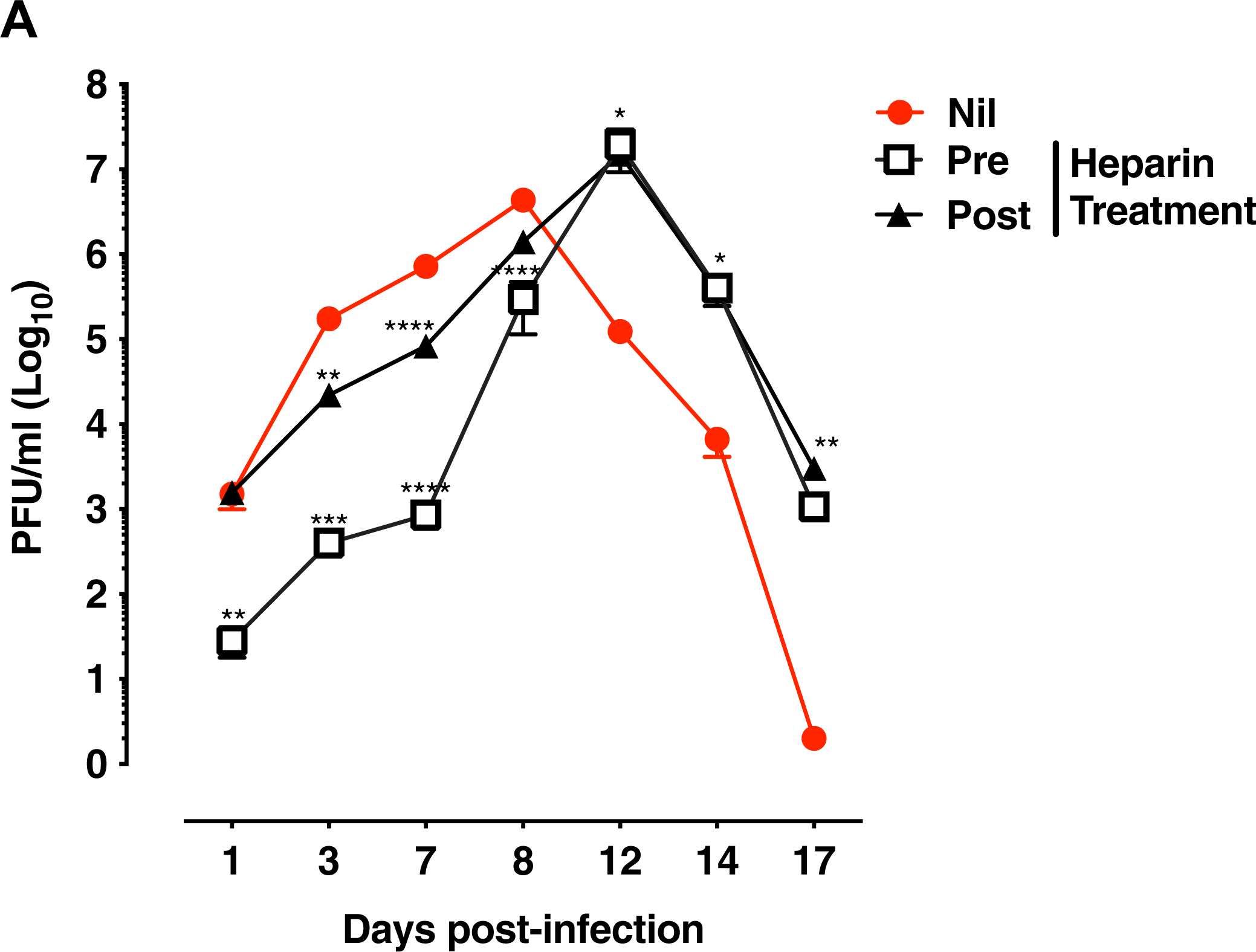
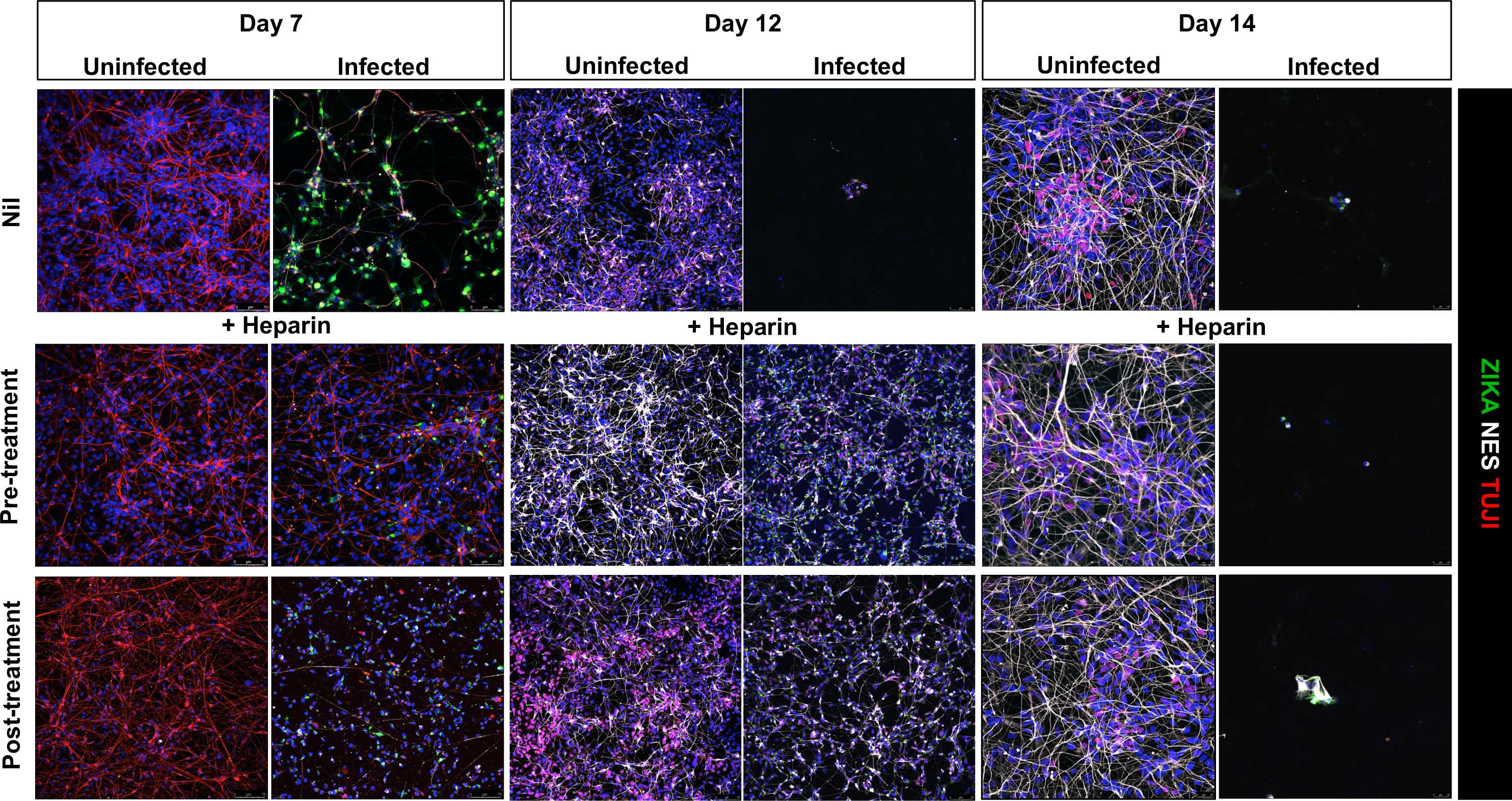
Kinetics of ZIKV replication in hiPSC-NPCs treated with a single dose of heparin. **A.** hiPSC-derived NPCs were treated with heparin (100 µg/ml) before and after infection (1h) with PRVABC59 isolate (MOI 0.001). Cell cultures were maintained in induction medium for 1 week, and then differentiation medium for at least 14 days. Kinetics of viral replication were determined by titering the infectious virus released in culture supernatants by PFA in Vero cells. Graphs represent the mean ± SD of 3 independent experiments. One-way Anova was used with Bonferroni correction. * Represents the statistical comparison among groups (**, p < 0.01; ***, p<0.001). **B.** Differentiated hiPSC-NPCs were fixed at day 7, 12 and 14 post-infection and stained for ZIKV with envelope protein (ZIKA-green); neurons with beta tubulin III (TUJ1-white) and PAX6 (red). Blue channel is DAPI staining for all cell types. Scale bar: 75 μm and 25 μm. Images were obtained using SP8 confocal microscope.

### Multiple heparin additions to hiPSC-NPCs during differentiation into neuro-glial cells induced a state of persistent ZIKV infection

As a single treatment of hiPSC-NPCs with heparin delayed ZIKV cytopathic effect but did not reverse it, heparin was added to the cultures twice a week up to 14 days. As shown in **Fig. 6A**, the kinetics of viral replication of heparin-treated cultures were delayed compared to those of untreated cultures; unlike a single heparin treatment, however, virus detection was extended in heparin-treated cells up to 17 days post-infection. Indeed, re-addition of heparin improved cell survival as shown in **Fig. 6B**. Nevertheless, by preventing virus-induced cell death, the differentiation of hiPSC-NPCs into neurons was preserved, despite ZIKV infection, as shown by the staining with a human mAb against envelope E protein (**Fig. 6B**).

**Fig. 6.**
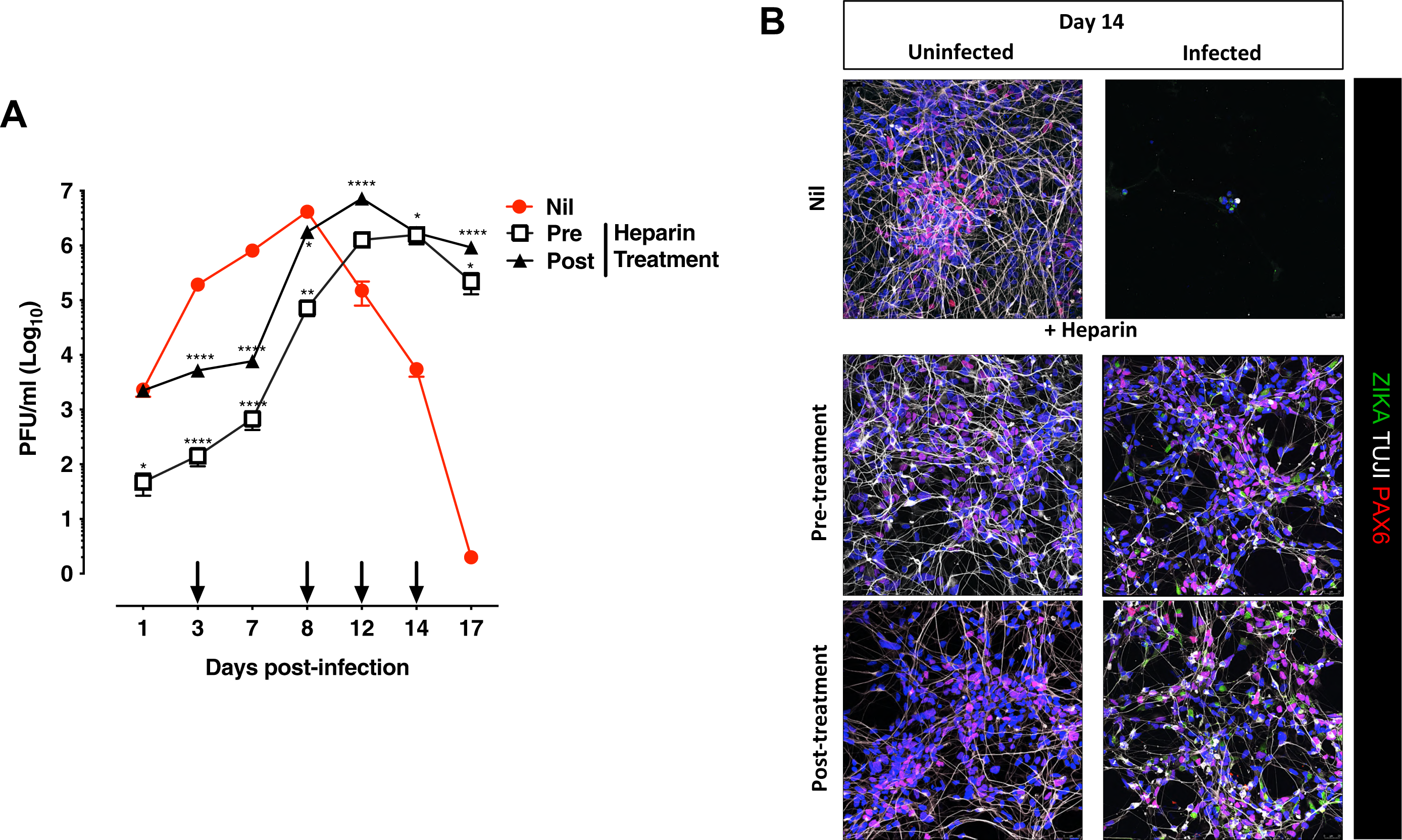
Kinetics of ZIKV replication in hiPSC-NPCs treated with multiple doses of heparin. **A.** HiPSC-derived NPCs were differentiated for 2 weeks. Cells were treated with heparin either before (pre) PRVABC59 infection (MOI 0.001) or after (post) infection. Heparin was added every 3 days (black arrows). Virion release in the supernatant was quantified at different time points by a plaque forming assay in Vero cells. The graph represents the mean ± SD of 3 independent experiments. One-way Anova with Bonferroni correction was used. * Represents statistical comparison among groups (**, p < 0.01; ***, p<0.001). **B.** Cells were fixed after 14 days since infection and stained for ZIKV with the anti-E antibody (ZIKA-green), neurons with beta tubulin III (TUJ1-white) and PAX6 (red). Blue channel is DAPI staining for all cell types. Scale bar: 75 μm. Images were obtained using SP8 confocal microscope.

### A single treatment of hf-NPCs with heparin prior to and post-infection preserves differentiation into neuro-glial cells

Unlike hiPSC-NPCs, hf-NPCs were induced to pan neuro-glia differentiation on a matrigel-coated surface and in NeuroCult™ NS-A Differentiation medium for at least 14 days. hf-NPCs were incubated either 1 h prior to or 4 h post-infection with heparin (100 µg/ml). Following viral inoculum removal, fresh differentiation medium was replaced twice a week for the following 2 weeks.

The kinetics of virus replication in cells incubated with heparin were similar to those of untreated cells (**Fig. 7A**); unlike hiPSC-NPCs a significant reduction of infectious virus released in the supernatant was observed in heparin-treated cultures at day 1 and 3 post-infection, especially in the pre-treatment condition. The proportion of ZIKV-positive cells increased at day 3 post-infection compared to day 1 in untreated cultures (**Fig. 7B**). Pre-incubation of cell cultures with heparin reduced the number of ZIKV-positive cells, whereas their exposure to heparin after infection reduced the number of infected cells at day 1 post-infection, although less efficiently than what was observed in the heparin pre-treatment condition. Cells were also stained with vimentin that is poorly expressed in embryonic stem cells and is switched on early in differentiation ^50^. As shown in **Fig. 7B**, in uninfected cells, vimentin expression increased after 3 days of culture. ZIKV infection in the untreated condition did not alter vimentin expression, though heparin post-treatment accelerated hf-NPC differentiation. Indeed, the differentiation status of the cells was examined after 7 days of differentiation (14 days post-infection) by staining the cell culture with TUJI and glial fibrillary acidic protein (GFAP), markers of neurons and astrocytes, respectively. As shown in **Fig. 7C**, uninfected cells were positive for GFAP and TUJI after 7 days post-differentiation, albeit the proportion of GFAP-positive cells was higher than TUJI-positive cells that became more visible 14 days post differentiation. At day 7, the GFAP signal was weak, and neurons were almost absent in infected untreated cells whereas infected hf-NPCs treated with heparin prior to infection differentiated into neurons (TUJI positive cells) and astrocytes (GFAP positive cells) distinct to untreated cells. At 14 days post-infection, neurons were well-represented in uninfected conditions, whereas ZIKV infection killed many cells, and the astrocytes that survived infection had a different morphology compared to those in uninfected conditions. As observed in **Fig. 7C**, a brilliant GFAP signal was present in infected untreated cells, a phenotype previously correlated with cell activation status during inflammatory insults ^51^. Heparin-treated astrocytes had an intermediate phenotype, suggesting that heparin might protect from inflammation insult.

**Fig. 7.**
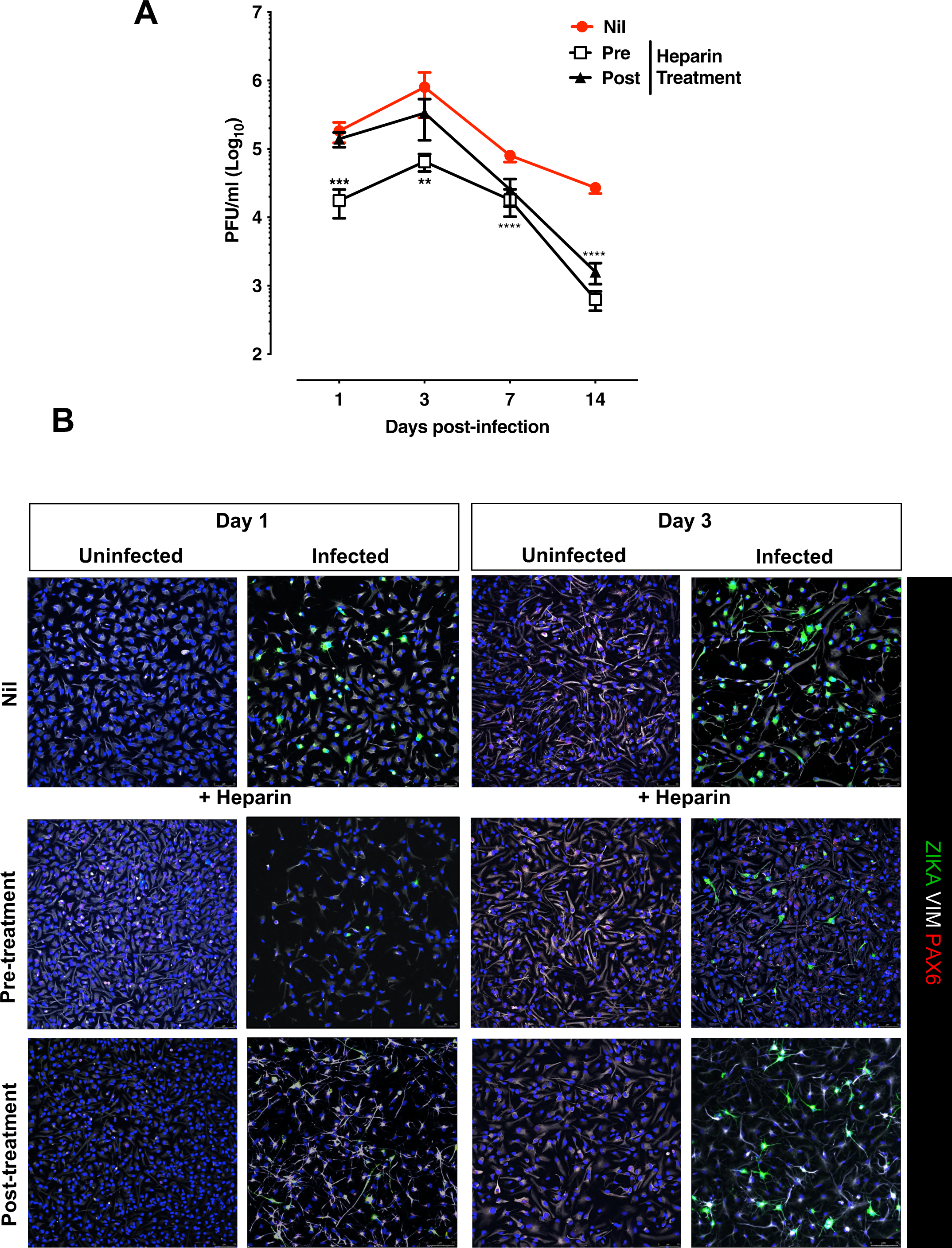
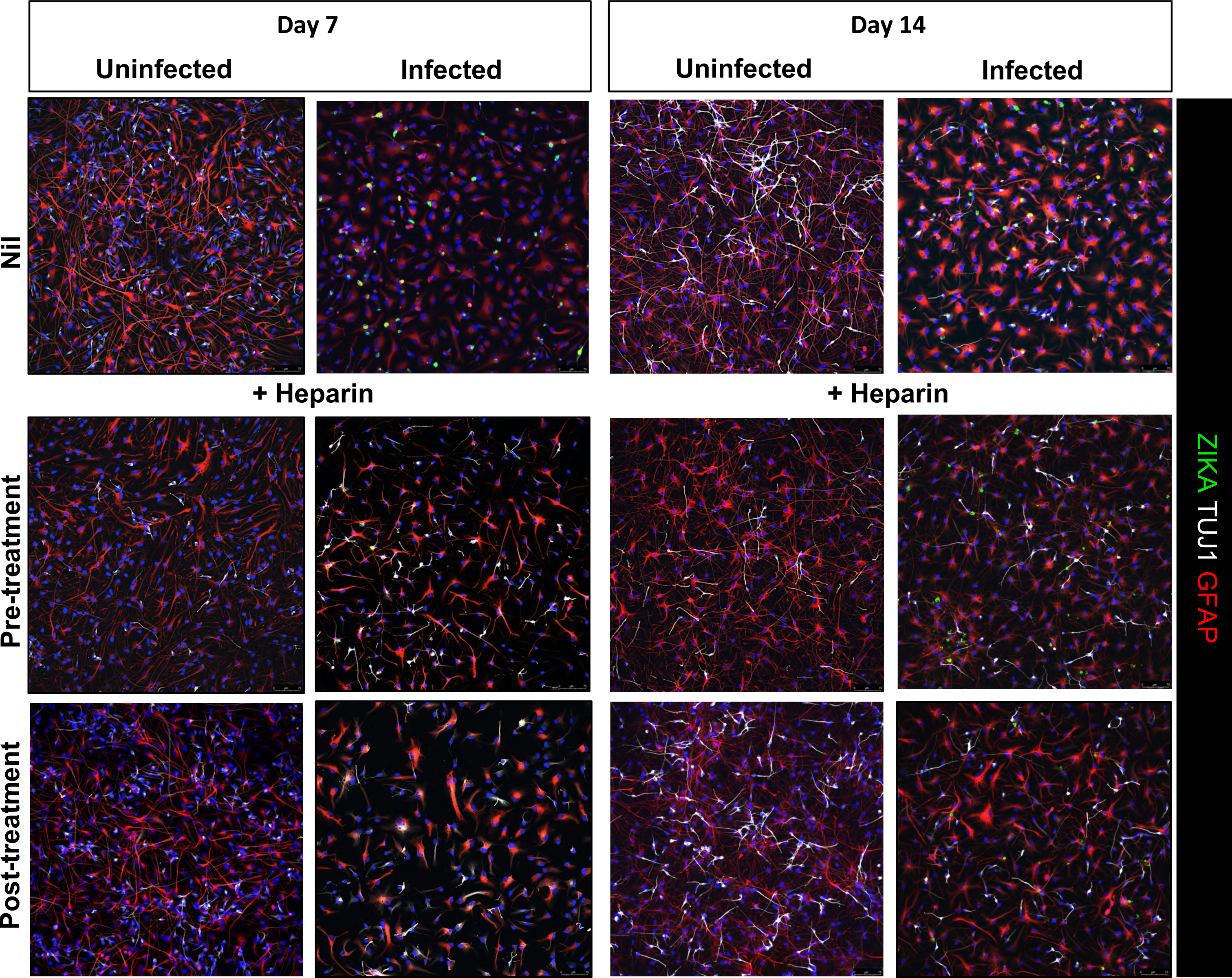
Kinetics of ZIKV replication in hf-NPCs treated with a single dose of heparin. **A.** Infectious virus released in supernatant was measured in a Vero-cell based Plaque Forming Assay. Cells were treated with heparin (100 µg/ml) either before (pre, 1 h) PRVABC59 infection (MOI 5) or after (post, for 4 h) infection. Cells were then differentiated for 2 weeks in a differentiation medium. The graph represents the mean ± SD of 3 independent experiments. One-way Anova with Bonferroni correction was used. * Represents statistical comparison among groups (**, p < 0.01; ***, p<0.001, ****, p<0.0001). **B.** Differentiated hf-NPCs were fixed at different time points; earlier time points (day 1 and 3) were stained for ZIKV with the anti-E antibody (ZIKA-green), Vimentin (white) and Pax6 (PAX6-red). Blue channel is DAPI staining for all cell types. Scale bar: 75 μm. Images were obtained using SP8 confocal microscope. **C.** Staining of cultures at later time points (day 7 and 14) post-infection as described in panel B. Scale bar: 75 μm. Images were obtained using a SP8 confocal microscope.

The percentage of neurons and astrocytes was quantified over the untreated uninfected cultures. As shown in **Fig. 8A**, a strong impairment in neuronal maturation was observed in the infected untreated condition (∼2% of neurons); the percentage of mature neural cells in infected cultures incubated with heparin was significantly higher than that observed in infected cultures (∼ 20% of neurons) and did not differ from the uninfected conditions (**Fig. 8A**). Astrocytes matured under all conditions although a significant decrease (∼15% of total cells) was observed in infected conditions compared to uninfected controls (∼40% of total cells); heparin partially restored glial maturation (∼30% of total cells) (**Fig. 8B**).

**Fig. 8.**
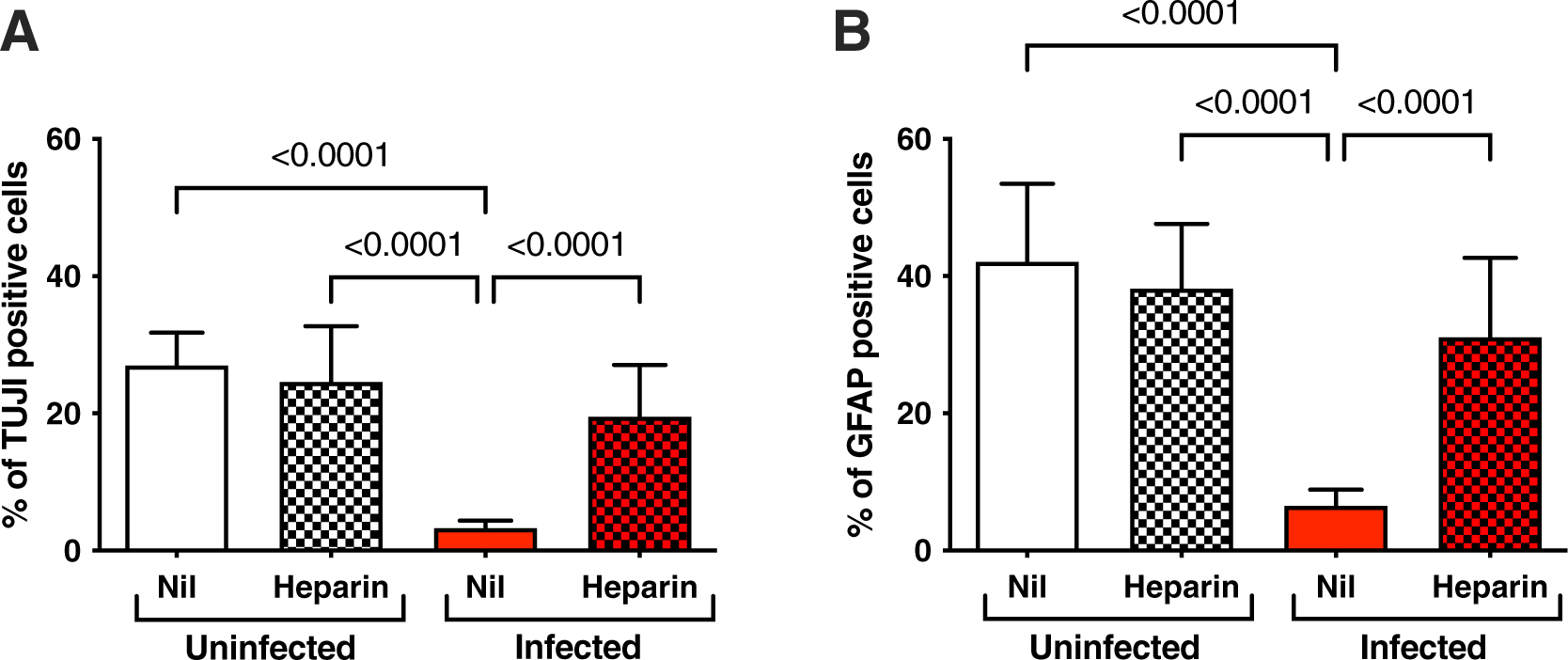
Quantification of neural-glial cells. **A.** Quantification of neurons. hf-NPCs were treated with heparin prior to infection with the PRVABC59-2015 isolate at the MOI of 5. After infection, cells were differentiated for 1 week. Cells were fixed after 7 days post-infection. **B.** Quantification of astrocytes. Three different images from 3 independent experiments were counted. Bars represent the percentage of neurons and astrocytes in the different conditions. P values were calculated by one-way Anova with the Bonferroni correction.

Similarly to hiPSC-NPCs, multiple heparin additions were tested in hf-NPCs either in pre-or post-treatment conditions. As shown in **Fig. 2SA**, these treatments resulted in a decrease of virion production 7 days post-infection with a concomitant increase of cell survival and differentiation vs. untreated conditions (**Fig. 2SB**).

## DISCUSSION

Heparin protects hNPCs grown in adherence or as neurospheres from ZIKV-induced apoptosis and non-apoptotic cell death, i.e. paraptosis. By inhibiting ZIKV-induced cell death, the release of infectious virions was diminished in heparin treated vs. untreated cultures. A single addition of heparin delayed ZIKV replication in hiPSC-NPCs committed to differentiate into neuro-glial cells. Multiple additions of heparin induced a prolonged infection with inhibition of virus-induced cell death and preserved neural-glial differentiation. Surprisingly, when differentiating hf-NPCs were infected in the presence of heparin as for hiPSC-NPCs, virus replication was significantly inhibited. In untreated cultures, virus persisted during their differentiation, while in heparin-treated cultures, ZIKV replication decreased, and hf-NPCs differentiated into neurons and glial cells. Overall, these results demonstrate the neuro-protective activity of heparin against the cytopathicity of ZIKV in hNPCs infection.

Similarly to other members of the Flaviviridae family, ZIKV causes cytopathic infection of fetal neural progenitor cells that leads to defective neurogenesis ^17, 52^ ^53^. A number of studies has reported apoptosis as a mechanism of cell death in ZIKV-infected hNPCs^54 28^, however, apoptosis has also been found in adjacent cells without evidence of infection since bystander cells are susceptible to the cytotoxic factors released during the cell death of productively infected cells ^16^. In addition to caspase-dependent cell death, a hallmark of apoptotic cell death, caspase-independent cytopathicity has also been reported in cells such as epithelial cells, primary skin fibroblasts and astrocytes ^33^. This mode of cell death was explained by massive vacuolization of the ER that is the major intracellular site of ZIKV replication ^33^. The accumulation of ZIKV vacuoles triggers a cellular collapse defined as paraptosis. Heparin treated hNPCs were protected by ZIKV-induced vacuolization, but also virus-induced apoptosis. This heparin activity was confirmed in hiPSC-NPCs and hf-NPC grown either in adherence or as neurospheres. As heparin does not transverse the plasma membrane, its protective mechanism must rely on molecular interactions at the cellular plasma level. Heparin is known to be a potent modulator of cellular receptors of growth factors such as fibroblast growth factor (FGF), epidermal growth factor (EGF) and vascular endothelial growth factor (VEGF) ^55^. Among the growth factors involved in neurogenesis, FGF-2, together with EGF, is essential for neuronal cell survival and maturation and plays a crucial role in CNS development ^56^. Heparin plays a critical role in the regulation of the FGF-2 activity by interacting with its receptor or stabilizing it and preventing its degradation. This interaction is specific to heparin and is not general to other proteoglycans ^57^.

Our previous work has shown that heparin prevents ZIKV-induced cell death without interfering with viral replication. In the present work, we have tested the protective effect of heparin on virus-induced cell death in the NS, a 3D system formed from spontaneous aggregations of hNPCs in non-adherent conditions ^18^. It is well established that the best 3D model for studying neurodevelopment is represented by brain organoids ^58^, however, their preparation from hNPCs is complex and costly. As alternative to brain organoids, NS have been shown to be permissive to ZIKV infection that disrupts their spherical structure by virus-induced cytopathic effects ^18^. We therefore tested whether heparin was effective in preventing cell death of ZIKV-infected NS derived from either hiPSC- or hf-NPCs. As expected, heparin treatment of NS strongly protected them from ZIKV-induced cytopathic effects by preventing NS disruption and inhibiting both apoptosis and necrosis. By maintaining NS integrity, heparin significantly inhibited the release of infectious virus in the culture supernatant. However, we cannot exclude that heparin might also exert antiviral effects. In our previous work, hNPCs were infected at a multiplicity of infection (MOI) of 1 ^40^, implying that on average, every cell was exposed to one virion. However, as the MOI could not be precisely calculated in the infection of NS as they are formed by cell aggregates, it is possible that NS were infected with a lower amount of infectious virus than NPCs grown in adherence. Alternatively, virions could only come into contact with cells present on the surface of the sphere. While cell-to-cell spreading infection is more efficient than cell-free infection ^59^, the amount of virions that reached the center of the NS is likely to be lower than that of a simple monolayer in which all cells are potentially exposed to the virus.

Although heparin significantly reduced vacuole formation, the number of ZIKV virions present in heparin treated cells was comparable to that of untreated infected controls. This finding raised the question of whether the cells maintained the ability to differentiate into neuro-glial cells. To check whether hNPCs treated with heparin are still functional, ZIKV infected hNPCs were differentiated into mature neural-glial cells. In hiPSC-NPCs, heparin delayed the kinetics of viral replication, protected hNPCs from ZIKV-induced cell death and allowed their differentiation into neurons and astrocytes. Interestingly, multiple additions of heparin induced prolonged viral replication. After the initial reduction of viral replication, virus spreading was similar in heparin treated cultures as compared to controls, while a proportion of cells survived and continued to produce virus. Specifically, we do not know whether the initial reduction of infected cells had segregated the virus to daughter cells during cell division and differentiation. Nevertheless, heparin is endowed with beneficial effects in promoting ZIKV-infected hNPCs differentiation into neuro-glial cells.

In addition to the hiPSC-NPC model, we also exploited the hf-NPCs because they recapitulate the heterogeneity of neural precursor cells of the embryonic human brain ^41^. Indeed, hf-NPCs were less susceptible to ZIKV cytopathic effect than hiPSC-NPCs and, after the initial cytopathic phase, a proportion of cells survived. By inhibiting ZIKV replication and preventing virus-induced cell death, heparin leaves the proportion of neurons and astrocytes comparable to that of uninfected cultures. The absence of neurons in infected culture could be due to inflammatory products induced by ZIKV infection that could influence cell maturation and was restored and maintained following cell incubation with heparin.

The distinct behaviour of hf-NPCs and hiPSC-NPCs could be explained by their diverse composition. Indeed, hiPSC-NPCs are homogeneous, whereas hf-NPCs consist of a heterogeneous mixture of stem cells and neuronal precursors at different early stages of differentiation ^60^. These features might resemble the *in vivo* neurodevelopment more closely than the hiPSC-derived NPCs, even though their limited availability and the heterogeneity of the source could be a major drawback. Apart from differences between these two cell culture systems, the protective activity of heparin against ZIKV-induced cell death is strongly preserved in all models used although further investigation is needed to determine the mechanism of this protection.

Collectively, these results highlight the potential of heparin protective activity against neurotropic viruses. Unfractionated heparin (UFH) as used here, is a highly sulphated, high molecular weight polysaccharide with anticoagulant activity. Given its high molecular weight, the proportion of UFH that can reach the fetus and protect the fetal brain from virus damage is likely low. Nevertheless, heparin can be chemically and enzymatically modified to obtain derivatives or fractions with low molecular weight, increased bioavailability and a more predictable anticoagulant activity and safety profile ^61^. Active fractions with low, or no anticoagulant activity may reside in crude heparin, the precursor material to pharmaceutical UFH, a resource awaiting further exploitation. In addition to its anticoagulant activity, heparin can prevent tissue injuries at the fetal–maternal interface ^62^. For instance, heparin improves successful embryo implantation by suppressing natural killer cell cytotoxicity ^63^, prevents leukocyte adhesion/influx ^64^, antagonizes interferon-γ (IFN-γ) signaling ^65^ and modulates chemokine activity ^66^. Mouse models of ZIKV infection, particularly pregnant mice will be instrumental to determine heparin beneficial effects to the fetal brain.

In conclusion, heparin and potentially its derivatives (devoid of anticoagulant activity), could play a role as antiviral agents, especially in providing rapid countermeasures against present and future emerging viral diseases, such as the current SARS-CoV-2 infection.

## MATERIALS AND METHODS

### Ethics statement

The study protocol was approved by the Ethical Committee of the IRCCS San Raffaele Hospital (Milan, Italy). Subjects participating in the study provided informed consent (Banca INSpe). The study conformed the standards of the Declaration of Helsinki.

### Human iPSC-derived NPCs (hiPSC-NPCs)

Fibroblasts were isolated from the skin biopsy of one healthy subject as described in ^67^. Fibroblasts were reprogrammed into iPSCs by using the episomal Sendai virus approach (CytoTune-iPS 1.0 Sendai Kit, Life Technologies) to obtain hiPSCs. Cells were maintained in feeder-free conditions in mTeSR1 culture medium (Stem Cell Technologies) on matrigel ES (Corning) coated plates and passaged using 0.5 mM EDTA. hNPCs were generated with some modification of the protocol described in Reinhardt *et al.,* ^68^. Briefly, colonies of hiPSCs grown on MaES (Corning) were detached using dispase (STEMCELL Technologies) and maintained in human embryonic stem cell (hESC) medium without bFGF2, supplemented with 1 µM dorsomorphin (Stemgent), 3 µM CHIR99021 (Tocris), 10 µM SB-431542 (Miltenyi) and 0.5 µM purmorphamine (Alexis). Embryoid bodies (EBs) were formed by culturing cells in non-culture Petri dishes (Greiner). On day 2, the medium was changed to N2B27 medium containing equal parts of neurobasal (Invitrogen) and DMEM-F12 medium (Invitrogen) with 1:100 B27 supplement lacking vitamin A (Invitrogen), 1:200 N2 supplement (Invitrogen), 1 % penicillin/streptomycin/glutamine (Gibco) and the same small molecules as used above. On day 4, dorsomorphin and SB-431542 were withdrawn, while 150 µM ascorbic acid was added to the medium. On day 6, EBs were mechanically dissociated into smaller aggregates and seeded onto matrigel (Matrigel Growth-factor-reduced, high concentration, Ma-GFRH Corning) coated 12-well plates (Corning). After hNPCs reached confluence, cells were detached with Accumax (Sigma) and replated (at least 1:5) in presence of ROCK inhibitor (Calbiochem). After 3 passages, purmorphamine was replaced by 1 µM SAG (Calbiochem). hNPCs were expanded until passage 10 before infection and oligo-dendroglial differentiation. NS were obtained by maintaining single cells in non-culture Petri dishes (Greiner) where they were grown as cell aggregates.

### Human fetal brain-isolated NPCs (hf-NPCs)

hf-NPCs are non-immortalized human fetal neural precursor cells (named BI-0194-008 cell line) obtained from a single human fetus 10-12 weeks post-conception (wpc) as reported in ^67^. Human tissue was provided by “Banca Italiana - Fondazione IRCCS CA’ GRANDA Ospedale Maggiore Policlinico di Milano”. Permission to use human fetal CNS tissue was granted by the ethical committee of the San Raffaele Hospital (approved on 13/06/2013). Tissue procurement was in agreement with the declaration of Helsinki and in agreement with the ethical guidelines of the European Network for Transplantation (NECTAR). The BI-0194-008 cell line is currently in use for the clinical trial EudraCT 2016-002020-86, NCT03269071. Briefly, primary, growth factor-expanded hNPCs were obtained as heterogeneous culture of spherical cell aggregates, derived from the diencephalic and telencephalic regions. Cells were grown in suspension as spheres in flask in NeuroCult-XF Proliferation Medium Human (STEMCELL Technologies) with EGF and bFGF (10 ng/ml each, R&D Systems). Every after 10-15 days, enzymatic dissociation of neurospheres with Accumax and re-plating (20-25000 cells/cm^2^) was performed.

### Differentiation of hNPCs

hiPSC-NPC-derived mature neuro-glial cells were generated by an induction phase followed by a differentiation step with modification of the protocol described in ^49^. Cells were seeded on Ma-GFR coated 12 mm diameter coverslips, and induction was allowed to occur in the presence of DMEM-f12 medium supplemented with N2 (1:50), B27 (1:50), MEM-NEAA (Gibco), PSG (Gibco), heparin (2 μg/ml), cAMP 1 μM (SIGMA), and 10 ng/ml insulin growth factor-1 (IGF-1). Half the medium was changed twice a week. After 7 days, the medium was replaced with neurobasal medium, N2 (1:50), B27 (1:50), MEM-NEAA (Gibco), PSG, supplemented with 1 μM cAMP, 150 µM AA, 10 ng/ml brain-derived neurotrophic factor (BDNF), 10 ng/ml IGF-1 and 10 ng/ml glial cell-derived neurotrophic factor (GDNF).

To obtain hf-NPCs pan-differentiation, spheres (passages 10-12) were dissociated in single cells and seeded on 12 mm (diameter) glass coverslips (70000/each) coated with matrigel GFR (0.2 μg/ml, Corning). Cells were maintained in NeuroCult™ NS-A Differentiation Medium Human (STEMCELL Technologies) for up to 14 days.

### Viruses

Three virus isolates were employed: the historical ZIKV strain (MR766), (EVAg - European Virus Archive), the Puerto Rico PRVABC59-2015 obtained from the CDC (GenBank Accession #KU501215) and the Brazilian 2016-INMI-1 (GenBank Accession # KU991811) obtained from an Italian individual who travelled to Brazil in January 2016. Viral isolates were expanded in Vero cells and titrated in a Plaque Forming Assay (PFA).

### ZIKV Infection and Ultrastructural Analysis

Adherent hiPSC-NPCs were infected at a MOI of 1 with the PRVABC59 isolate. Three days after infection, cells were fixed in 4% formaldehyde and 2.5% glutaraldehyde in cacodylate buffer and incubated for 5 min at room temperature. The samples were then fixed with 2% OsO4 in 2.5% glutaraldehyde in cacodylate buffer for 60 minutes. Monolayers were dehydrated in graded ethanol, washed in propylene oxide and infiltrated for 12 hours in a 1:1 mixture of propylene oxide and epoxide resin (Epon). Cells were then embedded in Epon and polymerized for 24 hours at 60°C. Slices were cut with an ultramicrotome (Ultracut Uct, Leica, Deerfield, IL), stained with uranyl acetate and lead citrate, and metaled. The ultrathin sections of infected hiPSC-NPCs were observed through transmission electron microscopy (Hitachi H7000).

### ZIKV Infection of NS

Both hiPSC- and hf-NPCs were grown spontaneously in rotation as NS after seeding for 3 days before infection. Heparin (Celsus Laboratories, Cincinnati, USA) was added 1 h prior to infection at a final concentration of 100 µg/ml. NS were infected with Brazilian 2016-INMI-1 isolates at a MOI of 1. Viral supernatants were collected 3 and 6 days post-infection and induced-cell death and viral titers were determined in the ToxiLight Bioassay (Lonza) and PFA, respectively. Brightfield images were captured at 3 and 6 days post-infection to measure NS diameter using ImageJ software. NS were transferred onto slides pre-coated with Matrigel and fixed at 3 and 6 days post-infection for the evaluation of the efficiency of infection by immunofluorescence with specific antibodies.

### ZIKV Infection of hNPCs induced to differentiate into neuro-glial cells

hiPSC-NPCs were seeded into 24-well plates at a final concentration of 100,000 cells per well. Twenty-four hours post-seeding, cells were infected with the PRVABC59 isolate at a MOI of 0.001 and treated with heparin (100 μg/ml). Two distinct protocols were adopted to test heparin effects on differentiation of hNPCs into neural-glial cells. In the pre-treatment protocol, cells were incubated with heparin 1 h prior to infection. After 4 h, the virus inoculum was removed and induction medium supplemented with cAMP and IGF-1 was added for 1 week, followed by replacement with differentiation medium for the following 3 weeks. In the post-treatment protocol, cells were infected for 4 h, then virus inoculum was removed and heparin was added for 1 h. Culture medium was replaced with the induction medium supplemented with cAMP and IGF-1 and changed every 3 days. After 1 week, the induction medium was replaced with the differentiation medium for the following 3 weeks. The differentiation medium was changed every 3 days. Supernatants were harvested at different time points to determine the infectious titers and cells were fixed side-by-side and cell differentiation during infection with specific antibodies.

hf-NPCs were placed in a 15 ml Falcon tube at a final concentration of 70,000 cells /ml. The PRVABC59 isolate was added at a MOI of 5 and treated with heparin (100 μg/ml). As described above, in the pre-treatment protocol, cells were incubated with heparin 1 h prior to infection. After 4 h, cells were centrifuged, resuspended in Neurocult™ NS-A differentiation medium (Stem Cell) and seeded into 24-well plates. Cultures were maintained for 14 days. In the post-treatment protocol, cells were infected for 4 h, cells were centrifuged, resuspended in Neurocult™ NS-A differentiation medium (Stem Cell) and heparin was added for 1 h. The differentiation medium was changed every 3 days up to 14 days. Supernatants were harvested at different time points to determine the infectious titers and cells were fixed side-by-side and cell differentiation during infection with specific antibodies.

### Immunofluorescence and image capture

NS were fixed using 4% paraformaldehyde (PFA) for 10 min, washed three times with PBS and incubated for 1 h with blocking solution PBS-0.5% Triton-X100, 5% donkey serum solution. NS were stained with mouse anti ds-RNA ZIKA, anti-rabbit cl-CASP3 and anti-Goat Vimentin overnight in blocking solution. After three washings with PBS-0.1% Triton-X100, Alexa Fluor-conjugated secondary antibodies were incubated for 2 hours in blocking solution at room temperature, washed three times with PBS-0.1% Triton-X100. DAPI (1 µg/mL; Sigma-Aldrich) in PBS was incubated for 10 minutes to counterstain the nuclei, washed three times and mounted in DAKO. NS and cells were visualized and acquired using a Leica SP8 confocal microscope (Alembic Facilities). Merged images were then generated using ImageJ software.

Differentiated hNPCs were fixed using 4% paraformaldehyde (PFA) for 10 min, washed three times with PBS and incubated for 1 h with blocking solution PBS-0.5% Triton-X100, 5% donkey serum solution. Cells were then stained with specific primary antibodies, in particular, mouse anti ds-RNA ZIKA, chicken anti-neuron-specific class III beta-tubulin (TuJ1, Biolegend) and anti-rabbit glial fibrillary acidic protein (GFAP, Dako) overnight in blocking solution. After three washings with PBS-0.1% Triton-X100, Alexa Fluor-conjugated secondary antibodies were incubated for 2 h in blocking solution at room temperature, washed three times with PBS-0.1% Triton-X100. DAPI (1 µg/mL; Sigma-Aldrich) in PBS was incubated for 10 minutes to counterstain the nuclei, washed three times and mounted in DAKO.

The human monoclonal antibody ZKA190-rIgG1 against the Flavivirus E protein was generated as previously described in ^69^. The mouse monoclonal J2 double-stranded RNA (1:300, mouse) was obtained from the English and Scientific Consulting Kft (Hungary). and cl-CASP3 (1:300, rabbit, 9661, Cell Signaling) were used. Anti-goat Alexa-fluor 633 (Invitrogen), anti-chicken Alexa-fluor 555 (Invitrogen), anti-rabbit Alexa-fluor 546 (Invitrogen) and anti-mouse Alexa-fluor 488 (Invitrogen) were used as secondary antibodies diluted in 1% BSA in PBS and maintained for 45 mins in the dark. Hoechst dye was used to stain the nuclei.

### Plaque Forming Assay (PFA)

Vero cells (1.2×10^6^) were seeded in 6-well culture plates. 24 h later, ten-fold dilutions of virus containing supernatants were prepared in culture medium supplemented with 1% heat-inactivated FBS and 1 mL of each dilution was added to the cells. The plates were incubated for 4 h at 37 °C. Unabsorbed virus was removed and 2 ml of culture medium supplemented with 1% methylcellulose (Sigma) were added to each well, followed by an incubation at 37 °C for 6 days. The methylcellulose overlay was removed and the cells were stained with 1% crystal violet in 70% methanol. Plaques were counted and expressed as plaque-forming units per mL (PFU/mL) ^32^.

### Cell death detection assay

ToxiLight^®^ BioAssay (Lonza) was used to measure the AK activity in culture supernatants as a marker of cell death. Briefly, 10 μl samples of culture supernatant were transferred on a half black 96 well plate (Costar). 50 μl of the detection reagent was added to each well and the plate was incubated for 10 min at room temperature. Luminescence was measured in a Mithras LB940 Microplate Reader (Berthold Technologies). The results were expressed as relative luminescent units (RLU).

### ELISA

The HMGB1 level was determined in sample supernatants with a commercially available ELISA kit (HMGB1 ELISA kit II; Shino-Test Corporation, Tokyo) according to the manufacturer’s protocol. Briefly, the wells of the microtiter strips are coated with purified anti-HMGB1 antibody. HMGB1 in the sample binds specifically to the immobilized antibody and is recognized by a second enzyme marked antibody. After substrate reaction, the HMGB1 concentration is determined by the colour intensity.

### Statistical analysis

Prism GraphPad software v. 8.0 (www.graphpad.com) was used for all statistical analyses. The Fisher’s exact test was used to compare the distribution of cells with and without vacuoles. Comparison between two groups was determined with the Student t test. Comparison among groups was performed using the one-way or two-way analysis of variance (Anova) and the Bonferroni’s multiple comparison test.

## Acknowledgments

Isabel Pagani conducted this study as partial fulfilment of her Ph.D. in Molecular Medicine, Program in Basic and Applied Immunology, International Ph.D. School, Vita-Salute San Raffaele University, Milan, Italy. We thank Alessandro Preti for technical assistance and Guido Poli for critical reading of the manuscript. This work was supported by funding from the Italian Ministry of Health (RF-2016-02364155).

## Author Contributions

I.P. and L.O. were involved in experimental design and execution, data analysis, and figure generation; P.P. performed electron microscopy analysis; S.G. expanded and titered viral stocks; E.B. cultured neural progenitor cells; D.C. provided human monoclonal antibody; M.E.B. provided the HMGB1 reagents and expertise; M.R.C. isolated and provided the viral strains; E.A.Y. provided heparin and valuable scientific discussion during data analysis and manuscript preparation; G.M. contributed with cells and expertise in the field of neural progenitor cells, guidance in experimental design and manuscript preparation; E.V. contributed expertise in the field of virology, guidance in experimental design and manuscript preparation.

## Supplementary Figures

**Fig. 1S.**
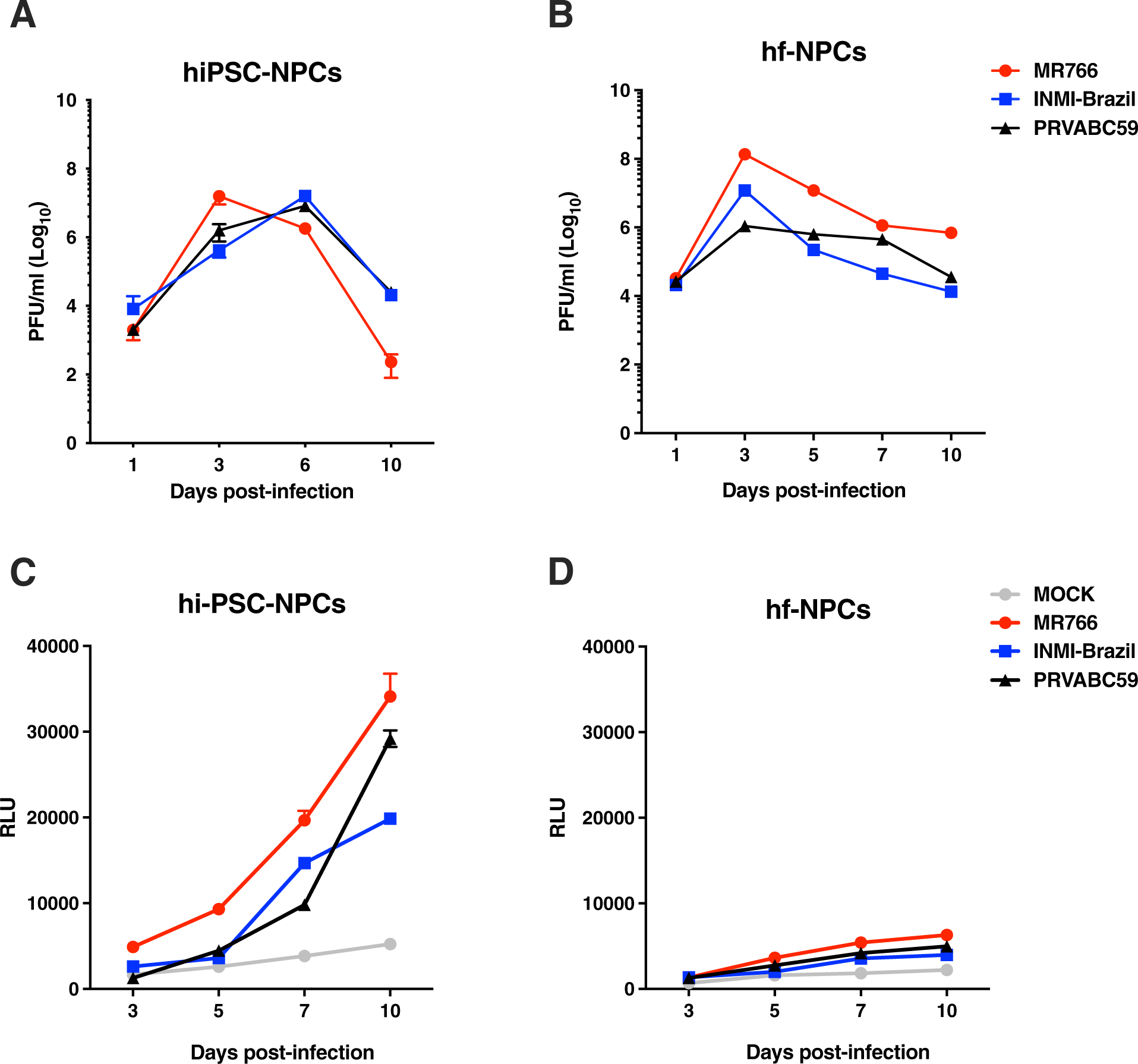
Kinetics of ZIKV replication and cytopathic effect in both hiPSC- and fetal-derived NPCs. hiPSC-(**A**) and hf-NPCs (**B**) were infected at a MOI of 1 with 3 different ZIKV strains: a historical MR766-1946, and two recent strains, i.e. PRVABC59 and Brazilian 2016-INMI-1. Supernatants were collected at 1, 3, 6 and 10 days post-infection. Infectious titers were determined by a plaque forming assay in Vero cells. The AK activity was measured in supernatants of infected hiPSC- (**C**) and hf-NPCs (**D**). The results are expressed as relative luminescent units (RLU). Means ± SD of two independent experiments are shown.

**Fig. S2.**
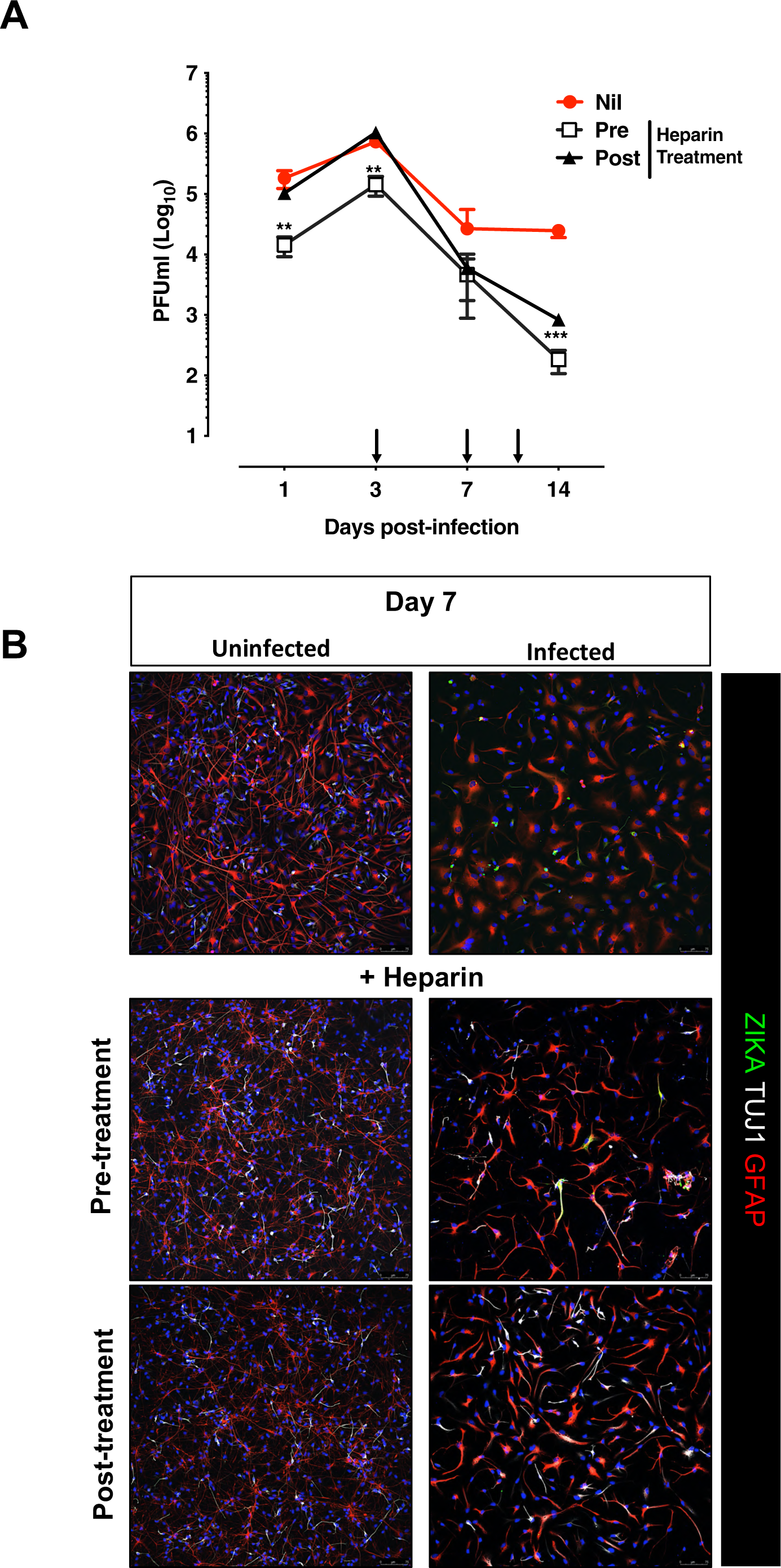
Kinetics of ZIKV infection after heparin multiple treatment in hf-NPCs during differentiation. **A.** Cells were treated with heparin (100 µg/ml) prior to and after infection (1 h) with the PRVABC59 isolate at a MOI of 5. The differentiation medium was added started 4 h post-infection. Heparin was added every 3 days up to 14 days post-infection. Infectious titers were measured in the culture supernatants by a Plaque Forming Assay in Vero cells. Nil is untreated infected cultures, pre is the treatment 1 h prior to infection; and post is the treatment 4 h post-infection. **B.** Cells were fixed after 7 days and stained for ZIKV with envelope protein (ZIKA-green); neurons with TUJI (white) and astrocytes with GFAP (red). Blue channel is DAPI staining for all cell types. Scale bar: 75 μm. Images were obtained using a SP8 confocal microscope.

## References

1. Musso D, Gubler DJ. Zika Virus. Clinical microbiology reviews 29, 487–524 (2016).

2. D’Ortenzio E, et al. Evidence of Sexual Transmission of Zika Virus. N Engl J Med 374, 2195–2198 (2016).

3. Moreira J, Lamas CC, Siqueira A. Sexual transmission of Zika virus: implications for clinical care and public health policy. Clinical infectious diseases : an official publication of the Infectious Diseases Society of America, (2016).

4. Cao-Lormeau VM, et al. Guillain-Barre Syndrome outbreak associated with Zika virus infection in French Polynesia: a case-control study. Lancet 387, 1531–1539 (2016).

5. Motta IJ, et al. Evidence for Transmission of Zika Virus by Platelet Transfusion. N Engl J Med 375, 1101–1103 (2016).

6. Dick GW. Zika virus. II. Pathogenicity and physical properties. Transactions of the Royal Society of Tropical Medicine and Hygiene 46, 521–534 (1952).

7. Haddow AD, et al. Genetic characterization of Zika virus strains: geographic expansion of the Asian lineage. PLoS neglected tropical diseases 6, e1477 (2012).

8. Duffy MR, et al. Zika virus outbreak on Yap Island, Federated States of Micronesia. N Engl J Med 360, 2536–2543 (2009).

9. Cao-Lormeau VM, et al. Zika virus, French polynesia, South pacific, 2013. Emerging infectious diseases 20, 1085–1086 (2014).

10. Brasil P, et al. Zika Virus Infection in Pregnant Women in Rio de Janeiro - Preliminary Report. The New England journal of medicine, (2016).

11. Driggers RW, et al. Zika Virus Infection with Prolonged Maternal Viremia and Fetal Brain Abnormalities. The New England Journal of Medicine 374, 2142–2151 (2016).

12. Hazin AN, et al. Computed Tomographic Findings in Microcephaly Associated with Zika Virus. The New England journal of medicine, (2016).

13. Besnard M, et al. Congenital cerebral malformations and dysfunction in fetuses and newborns following the 2013 to 2014 Zika virus epidemic in French Polynesia. Euro surveillance : bulletin Europeen sur les maladies transmissibles = European communicable disease bulletin 21, (2016).

14. Calvet G, et al. Detection and sequencing of Zika virus from amniotic fluid of fetuses with microcephaly in Brazil: a case study. The Lancet Infectious diseases, (2016).

15. Mlakar J, et al. Zika Virus Associated with Microcephaly. The New England journal of medicine 374, 951–958 (2016).

16. Ho CY, et al. Differential neuronal susceptibility and apoptosis in congenital Zika virus infection. Ann Neurol 82, 121–127 (2017).

17. Tang H, et al. Zika Virus Infects Human Cortical Neural Progenitors and Attenuates Their Growth. Cell stem cell 18, 587–590 (2016).

18. Garcez PP, et al. Zika virus impairs growth in human neurospheres and brain organoids. *Science*, (2016).

19. Souza BS, et al. Zika virus infection induces mitosis abnormalities and apoptotic cell death of human neural progenitor cells. Sci Rep 6, 39775 (2016).

20. Onorati M, et al. Zika Virus Disrupts Phospho-TBK1 Localization and Mitosis in Human Neuroepithelial Stem Cells and Radial Glia. Cell Rep 16, 2576–2592 (2016).

21. Nowakowski TJ, Pollen AA, Di Lullo E, Sandoval-Espinosa C, Bershteyn M, Kriegstein AR. Expression Analysis Highlights AXL as a Candidate Zika Virus Entry Receptor in Neural Stem Cells. Cell Stem Cell 18, 591–596 (2016).

22. Dang J, et al. Zika Virus Depletes Neural Progenitors in Human Cerebral Organoids through Activation of the Innate Immune Receptor TLR3. Cell Stem Cell 19, 258–265 (2016).

23. Hanners NW, et al. Western Zika Virus in Human Fetal Neural Progenitors Persists Long Term with Partial Cytopathic and Limited Immunogenic Effects. Cell Rep 15, 2315–2322 (2016).

24. Yoon KJ, et al. Zika-Virus-Encoded NS2A Disrupts Mammalian Cortical Neurogenesis by Degrading Adherens Junction Proteins. Cell Stem Cell 21, 349–358 e346 (2017).

25. Morrison TE, Diamond MS. Animal Models of Zika Virus Infection, Pathogenesis, and Immunity. J Virol 91, (2017).

26. Miner JJ, et al. Zika Virus Infection during Pregnancy in Mice Causes Placental Damage and Fetal Demise. Cell 165, 1081–1091 (2016).

27. Cugola FR, et al. The Brazilian Zika virus strain causes birth defects in experimental models. Nature 534, 267–271 (2016).

28. Hamel R, et al. Biology of Zika Virus Infection in Human Skin Cells. Journal of virology 89, 8880–8896 (2015).

29. Vicenzi E, et al. Subverting the mechanisms of cell death: flavivirus manipulation of host cell responses to infection. Biochem Soc Trans 46, 609–617 (2018).

30. Cortese M, et al. Ultrastructural Characterization of Zika Virus Replication Factories. Cell Rep 18, 2113–2123 (2017).

31. Gladwyn-Ng I, et al. Stress-induced unfolded protein response contributes to Zika virus-associated microcephaly. Nat Neurosci 21, 63–71 (2018).

32. Volpi VG, Pagani I, Ghezzi S, Iannacone M, D’Antonio M, Vicenzi E. Zika Virus Replication in Dorsal Root Ganglia Explants from Interferon Receptor1 Knockout Mice Causes Myelin Degeneration. Sci Rep 8, 10166 (2018).

33. Monel B, et al. Zika virus induces massive cytoplasmic vacuolization and paraptosis-like death in infected cells. EMBO J 36, 1653–1668 (2017).

34. Sperandio S, Poksay KS, Schilling B, Crippen D, Gibson BW, Bredesen DE. Identification of new modulators and protein alterations in non-apoptotic programmed cell death. J Cell Biochem 111, 1401–1412 (2010).

35. Luo Y, et al. High mobility group box 1 released from necrotic cells enhances regrowth and metastasis of cancer cells that have survived chemotherapy. Eur J Cancer 49, 741–751 (2013).

36. Scaffidi P, Misteli T, Bianchi ME. Release of chromatin protein HMGB1 by necrotic cells triggers inflammation. Nature 418, 191–195 (2002).

37. McCauley MJ, Zimmerman J, Maher LJ, 3rd, Williams MC. HMGB binding to DNA: single and double box motifs. J Mol Biol 374, 993–1004 (2007).

38. Lange SS, Mitchell DL, Vasquez KM. High mobility group protein B1 enhances DNA repair and chromatin modification after DNA damage. Proc Natl Acad Sci U S A 105, 10320–10325 (2008).

39. Bianchi ME, Crippa MP, Manfredi AA, Mezzapelle R, Rovere Querini P, Venereau E. High-mobility group box 1 protein orchestrates responses to tissue damage via inflammation, innate and adaptive immunity, and tissue repair. Immunol Rev 280, 74–82 (2017).

40. Ghezzi S, et al. Heparin prevents Zika virus induced-cytopathic effects in human neural progenitor cells. Antiviral Res 140, 13–17 (2017).

41. Stein JL, et al. A quantitative framework to evaluate modeling of cortical development by neural stem cells. Neuron 83, 69–86 (2014).

42. Kobolak J, et al. Human Induced Pluripotent Stem Cell-Derived 3D-Neurospheres are Suitable for Neurotoxicity Screening. Cells 9, (2020).

43. Dmitriev RI, Zhdanov AV, Nolan YM, Papkovsky DB. Imaging of neurosphere oxygenation with phosphorescent probes. Biomaterials 34, 9307–9317 (2013).

44. Vicenzi E, et al. Coronaviridae and SARS-associated coronavirus strain HSR1. Emerg Infect Dis 10, 413–418 (2004).

45. Mycroft-West CJ, et al. Heparin Inhibits Cellular Invasion by SARS-CoV-2: Structural Dependence of the Interaction of the Spike S1 Receptor-Binding Domain with Heparin. Thromb Haemost 120, 1700–1715 (2020).

46. Noctor SC, Martinez-Cerdeno V, Kriegstein AR. Distinct behaviors of neural stem and progenitor cells underlie cortical neurogenesis. J Comp Neurol 508, 28–44 (2008).

47. Morshead CM, et al. Neural stem cells in the adult mammalian forebrain: a relatively quiescent subpopulation of subependymal cells. Neuron 13, 1071–1082 (1994).

48. Morest DK, Silver J. Precursors of neurons, neuroglia, and ependymal cells in the CNS: what are they? Where are they from? How do they get where they are going? Glia 43, 6–18 (2003).

49. Muratore CR, Srikanth P, Callahan DG, Young-Pearse TL. Comparison and optimization of hiPSC forebrain cortical differentiation protocols. PLoS One 9, e105807 (2014).

50. Chen M, et al. Increased Neuronal Differentiation of Neural Progenitor Cells Derived from Phosphovimentin-Deficient Mice. Mol Neurobiol 55, 5478–5489 (2018).

51. Ledur PF, et al. Zika virus infection leads to mitochondrial failure, oxidative stress and DNA damage in human iPSC-derived astrocytes. Sci Rep 10, 1218 (2020).

52. Rasmussen SA, Jamieson DJ, Honein MA, Petersen LR. Zika Virus and Birth Defects--Reviewing the Evidence for Causality. N Engl J Med 374, 1981–1987 (2016).

53. Turrini F, Ghezzi S, Pagani I, Poli G, Vicenzi E. Zika Virus: a re-emerging pathogen with rapidly evolving public health implications. New Microbiol 39, 86–90 (2016).

54. Wen Z, Song H, Ming GL. How does Zika virus cause microcephaly? Genes Dev 31, 849–861 (2017).

55. Li Y, Sun C, Yates EA, Jiang C, Wilkinson MC, Fernig DG. Heparin binding preference and structures in the fibroblast growth factor family parallel their evolutionary diversification. Open Biol 6, (2016).

56. Raballo R, Rhee J, Lyn-Cook R, Leckman JF, Schwartz ML, Vaccarino FM. Basic fibroblast growth factor (Fgf2) is necessary for cell proliferation and neurogenesis in the developing cerebral cortex. J Neurosci 20, 5012–5023 (2000).

57. Caldwell MA, Svendsen CN. Heparin, but not other proteoglycans potentiates the mitogenic effects of FGF-2 on mesencephalic precursor cells. Exp Neurol 152, 1–10 (1998).

58. Qian X, Song H, Ming GL. Brain organoids: advances, applications and challenges. Development 146, (2019).

59. Clark AE, Zhu Z, Krach F, Rich JN, Yeo GW, Spector DH. Zika virus is transmitted in neural progenitor cells via cell-to-cell spread and infection is inhibited by the autophagy inducer trehalose. J Virol, (2020).

60. Ottoboni L, von Wunster B, Martino G. Therapeutic Plasticity of Neural Stem Cells. Front Neurol 11, 148 (2020).

61. Greer I, Hunt BJ. Low molecular weight heparin in pregnancy: current issues. Br J Haematol 128, 593–601 (2005).

62. Hills FA, et al. Heparin prevents programmed cell death in human trophoblast. Mol Hum Reprod 12, 237–243 (2006).

63. Kumar P, Mahajan S. Preimplantation and postimplantation therapy for the treatment of reproductive failure. J Hum Reprod Sci 6, 88–92 (2013).

64. Wan JG, Mu JS, Zhu HS, Geng JG. N-desulfated non-anticoagulant heparin inhibits leukocyte adhesion and transmigration in vitro and attenuates acute peritonitis and ischemia and reperfusion injury in vivo. Inflamm Res 51, 435–443 (2002).

65. Fritchley SJ, Kirby JA, Ali S. The antagonism of interferon-gamma (IFN-gamma) by heparin: examination of the blockade of class II MHC antigen and heat shock protein-70 expression. Clin Exp Immunol 120, 247–252 (2000).

66. Spratte J, Schonborn M, Treder N, Bornkessel F, Zygmunt M, Fluhr H. Heparin modulates chemokines in human endometrial stromal cells by interaction with tumor necrosis factor alpha and thrombin. Fertil Steril 103, 1363–1369 (2015).

67. Butti E, et al. Neural precursor cells contribute to decision-making by tuning striatal connectivity via secretion of IGFBPL-1. bioRxiv DOI: 10.1101/2020.12.29.424678, (2020).

68. Reinhardt P, et al. Derivation and expansion using only small molecules of human neural progenitors for neurodegenerative disease modeling. PLoS One 8, e59252 (2013).

69. Wang J, et al. A Human Bi-specific Antibody against Zika Virus with High Therapeutic Potential. Cell 171, 229–241 e215 (2017).

